# Replicative Aging Remodels Cell Wall and is Associated with Increased Intracellular Trafficking in Human Pathogenic Yeasts

**DOI:** 10.1101/2022.01.25.477803

**Authors:** Vanessa K.A. Silva, Somanon Bhattacharya, Natalia Kronbauer Oliveira, Anne G. Savitt, Daniel Zamith-Miranda, Joshua D. Nosanchuk, Bettina C. Fries

**Author notes:** Address correspondence to Bettina C. Fries,. These authors have contributed equally to this work (Vanessa K.A. Silva, Somanon Bhattacharya).

## Abstract

Replicative aging is an underexplored field of research in medical mycology. *Cryptococcus neoformans* (*Cn*) and *Candida glabrata* (*Cg*) are dreaded fungal pathogens that cause fatal invasive infections. The fungal cell wall is essential for yeast viability and pathogenesis. In this study, we provide data characterizing age-associated modifications to the cell wall of *Cn* and *Cg*. Here, we report that old yeast cells upregulate genes of cell wall biosynthesis, leading to cell wall reorganization, and increased levels of all major components, including glucan, chitin and its derivatives, as well as mannan. This results in a significant thickening of the cell wall in aged cells. Old generation yeast cells exhibited drastic ultrastructural changes, including the presence of abundant vesicle-like particles in the cytoplasm, and enlarged vacuoles with altered pH homeostasis. Our findings suggest that the cell wall modifications could be enabled by augmented intracellular trafficking. This work furthers our understanding of the cell phenotype that emerges during aging. It highlights differences in these two fungal pathogens and elucidates mechanisms that explain the enhanced resistance of old cells to antifungals and phagocytic attacks.

**IMPORTANCE:** *Cryptococcus neoformans* and *Candida glabrata* are two opportunistic human fungal pathogens that cause life-threatening diseases. During infection, both microorganisms have the ability to persist for long periods, and treatment failure can occur even if standard testing identifies the yeasts to be sensitive to antifungals. Replicative lifespan is a trait that is measured by the number of divisions a cell undergoes before death. Aging in fungi is associated with enhanced tolerance to antifungals and resistance to phagocytosis, and characterization of old cells may help identify novel antifungal targets. The cell wall remains an attractive target for new therapies because it is essential for fungi and is not present in humans. This study shows that the organization of the fungal cell wall changes remarkably during aging and becomes thicker and is associated with increased intracellular trafficking as well as the alteration of vacuole morphology and pH homeostasis.

## INTRODUCTION

Fungal pathogens are an emerging threat to global health due to the rise of immunosuppressed patients (1). Their prevalence may even be affected by the evolving climate change (2, 3). Among the human fungal pathogens, *Candida* and *Cryptococcus* account for most of the invasive fungal infections (4). *Cryptococcus neoformans* (*Cn*), a member of the phylum Basidiomycota, is the leading agent of fungal meningitis and affects mostly patients living with advanced HIV disease (5). *Candida glabrata* (*Cg*), a member of the phylum Ascomycota, is the second most common fungal pathogen that causes bloodstream infections in North America and is associated with rapidly acquired antifungal resistance (6, 7). Currently, there are no vaccines for fungal diseases, and antifungal therapeutic options are still limited (8, 9).

The fungal cell wall is an ideal target for antifungal chemotherapies since it is essential for fungal viability and virulence, but absent in human cells (10). The fungal cell wall is composed of a complex matrix of glucose, proteins, lipids, and pigments (11). The cell wall of *Cn* is mainly composed of polymers of glucose (α and β -glucans), N-acetyl-glucosamine (GlcNAc) (chitin and chitooligomers), glucosamine (chitosan), and glycoproteins (mannoproteins) (12). Similarly, the cell wall of *Cg* is mainly composed of α and β -glucans, mannoproteins, and chitin (10).

Glucans and chitin are the most important structural components of fungal cell walls (13). Chitin can be degraded to chitooligomers, chitin-derived structures composed of 3-20 residues of GlcNAc (14). Chitooligomers are found around nascent buds and participate in connecting the cryptococcal cell wall to the capsule (14, 15). Also, chitin can be enzymatically deacetylated to chitosan (16). Most fungal pathogens expose chitin, but *Cn* is one of the few species that can evade the host immune system by replacing the chitin in its cell walls with chitosan (17). Mannose residues by N- or O- linkages can be found in association with proteins and are highly immunogenic (11).

The fungal cell wall is a dynamic structure that regulates cellular morphology (18), and remodels its components in response to environmental stresses, morphogenesis, cellular growth, and cell division (19). Aging is the consequence of cell division and a conserved natural process among eukaryotic cells. Replicative aging is the result of asymmetric cell divisions, and is an important contributor to pathogenesis (20). During replicative aging, the fungal cell wall becomes thicker, which is associated with increased resistance to antifungals and phagocytic uptake (21, 22). However, the changes in cell wall composition and architecture during aging have not been elucidated. In this study, we investigate cell wall remodeling during aging in *Cn* and *Cg*. We report the increased expression of essential genes related to fungal cell wall biosynthesis. These transcriptome changes concur with our observation of enhanced levels of significant cell wall components (chitin, chitooligomers and glucans) and enhanced cell wall thickness. Cell wall remodeling during aging is accompanied by formation of intracellular vesicles and multivesicular bodies, which suggests intensification of vesicular trafficking. In addition, our data indicate modifications in the vacuole morphology and pH, underscoring the importance of this organelle in the dynamics of progressive aging. Our data provide insights into the complex aging-related changes in fungal pathogens.

## RESULTS

### Regulation of cell wall biosynthesis genes is altered in older generation cells

Old yeast cells previously biotin-labeled and conjugated to streptavidin were separated when reached the desired generation using a magnetic field, while non-labeled young cells were washed off from the system. After isolation, we first performed a focused transcriptome analysis on known essential cell wall biosynthesis genes. Of the eight chitin synthases in *Cn* and four in *Cg*, *CHS3* generates the majority of chitin (16, 23). *CHS3* was upregulated in old *Cn* (3.3 fold-change, p ≤ 0.0001) (Fig. 1A) but not in old *Cg* (Fig. 1B). Likewise, we observed upregulations of *CHI22,* an endochitinase involved in chitooligomer formation (24) in older *Cn* (2.38 fold-change, p ≤ 0.0001) (Fig.1A) and *CTS1*, an endochitinase (2.66 fold-change, p ≤ 0.0001) in older *Cg* (Fig.1B).

**FIG 1.**
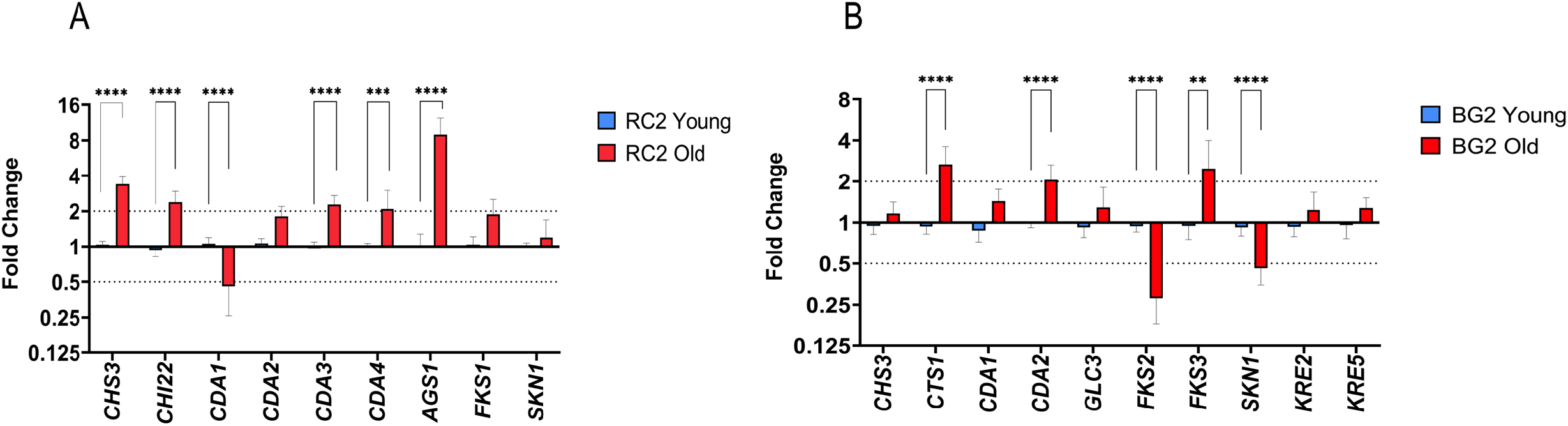
Increased expression levels of genes related to cell wall biosynthesis in old *C. neoformans* (*Cn*) and *C. glabrata* (*Cg*). Gene expression was analyzed by qRT-PCR in young (blue bars) and old (red bars) cells of RC2 *Cn* (A) and BG2 *Cg* (B). The data were normalized according to the gene expression of young cells, and the gene actin 1 (*ACT1*) was used as an internal control. The dotted lines represent two-fold up or down of the respective genes. All experiments were done in technical triplicates and repeated at least two times independently. Error bars correspond to the standard deviations of at least two independent experiments (n=2). *t*-test was performed to determine the *p*-value (**, p ≤ 0.01; ***, p ≤ 0.001; ****, p ≤ 0.0001).

In *Cn*, chitin can be converted to chitosan by four putative chitin deacetylases (*CDAs*) (17, 25). We found *CDA1* trancription was downregulated (0.45 fold-change, p ≤ 0.0001) whereas both *CDA3* and *CDA4* were upregulated (2.27 fold-change, and 2.08 fold-change, p ≤ 0.001, respectively) in old *Cn* (Fig.1A). In *Candida* species, chitosan is present only in the chlamydospore cell wall and is generated by two genes (*CDA1* and *CDA2*) (26), of which *CDA2* was upregulated (>2-fold-change, p ≤ 0.0001) in old *Cg* (Fig.1B). This is different from the *Cn*, where *CDA2* was not upregulated in the old cells (Fig. 1B).

Transcription of *AGS1*, which encodes for an α-1,3-glucan synthase, was upregulated in old *Cn* (9-fold-change, p ≤ 0.0001) (Fig.1A). In contrast, transcription of α-1,4-glucan synthase (*GLC3*) of *Cg* was not significantly altered Fig.1B). Genes relevant for β-glucan synthesis, *FKS1* and *SKN1,* did not significantly change in old *Cn* (Fig.1A) but were down-regulated in old *Cg*, genes *FKS2* (0.27 fold-change, p ≤ 0.05) and *SKN1* (0.46 fold-change, p ≤ 0.0001). *FKS3* was increased (2.48-fold-change, p ≤ 0.0001), whereas *KRE2* and *KRE5* remained not markedly altered in older *Cg* (Fig.1B).

### Yeast cell wall architecture is reshaped during aging

Next, we assessed the cell wall main components by fluorescent microscopy (Fig.2). In old *Cn*, chitin staining (by Calcofluor White, CFW) was enhanced surrounding the cell surface compared to younger cells (Fig. 2A). In old *Cg*, chitin staining was enhanced but remained localized to bud scars in the cell wall (Fig. 2B). A similar pattern was noted with the binding of wheat germ lectin (WGA), which binds to chitooligomers. Again, it was abundant throughout the cell wall of most old *Cn* cells. In young *Cn*, chitooligomers are located in the area of the emerging developing buds (Fig.2C). The latter pattern was similar to chitooligomer staining in young *Cg.* WGA staining of old *Cg* is also bound only to the numerous persistent bud scars (Fig.2D), similar to the calcofluor binding to chitin. The deacetylated form of chitin, chitosan, binds specifically to the anionic dye Eosin Y (25). Chitosan is uniformly enhanced and distributed through old *Cn* cell walls (Fig.2E). In contrast, chitosan was not detected in *Cg* cell walls (data not shown).

**FIG 2.**
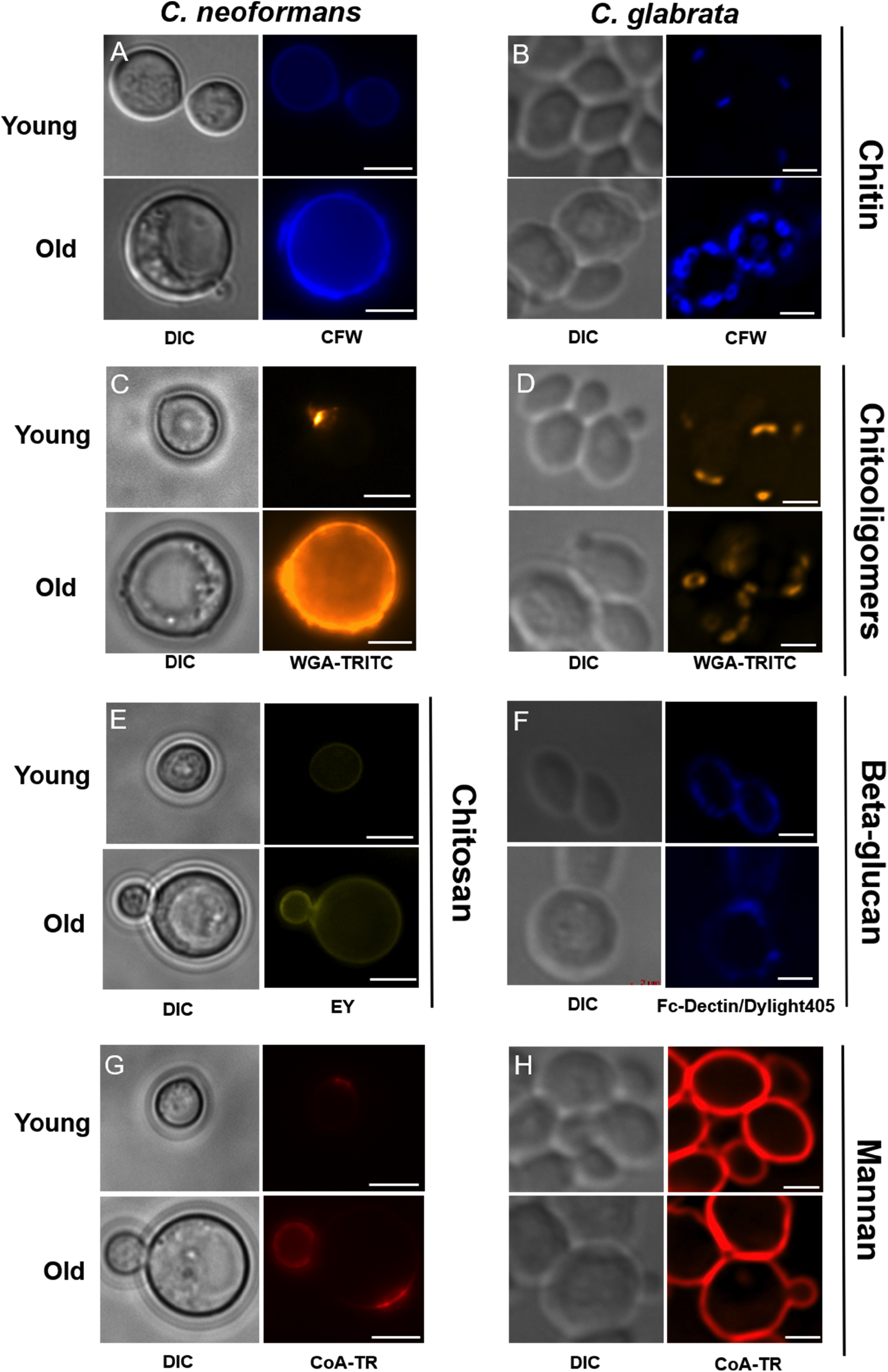
Cell wall architecture of *C. neoformans* (*Cn*) and *C. glabrata* (*Cg*) is remodeled during aging. Total chitin (blue) was stained with calcofluor white (CFW) (A and B), exposed chitooligomers (orange) with tetramethylrhodamine-labeled wheat germ agglutinin (WGA-TRITC) (C and D), chitosan (green) with Eosin Y (EY) (E), β-glucan (blue) with Fc-Dectin 1/Dylight 405 goat anti-human IgG + IgM (F), mannan (red) with concanavalin-Texas red (CoA-TR) (G-H). Stained cells were image by fluorescent microscopy, using the following channels: DAPI (CFW), DsRed (WGA-TRITC and CoA-TR) and GFP (EY). DIC, differential interference contrast. Imaging of at least 100 cells of *Cn* (RC2) and *Cg* (BG2) was performed at 100x in a Zeiss Axio Observer microscope, and a Zeiss Axiovert 200M microscope (Thornwood, NY), respectively. Images were processed using ImageJ/Fiji software. Scale bars correspond to 5 μm for *Cn*, and 2 μm for *Cg*.

The *Cn* polysaccharide capsule prevents binding of mAbs MOPC-104E (α-glucan) and Fc-Dectin (β-glucan), hence, like others, we were not able to assess α- and β-glucan levels (27, 28). Both young and old *Cg* exhibited comparable β-glucan staining of the cell surface (Fig.2F). Finally, we found that mannoproteins were enriched in the region next to the developing buds in both young and old *Cn*. However, old *Cn* presented slightly greater intensity than young cells (Fig.2G). Furthermore, mannoproteins of old cells also displayed a dotted staining pattern and co-localized in the capsule (Fig.3B), which was not observed in young cells (Fig.3A). In *Cg*, continuous staining of mannoproteins on the outer surface of the cell wall of young and old *Cg* cells was comparable (Fig.2H).

**FIG 3.**
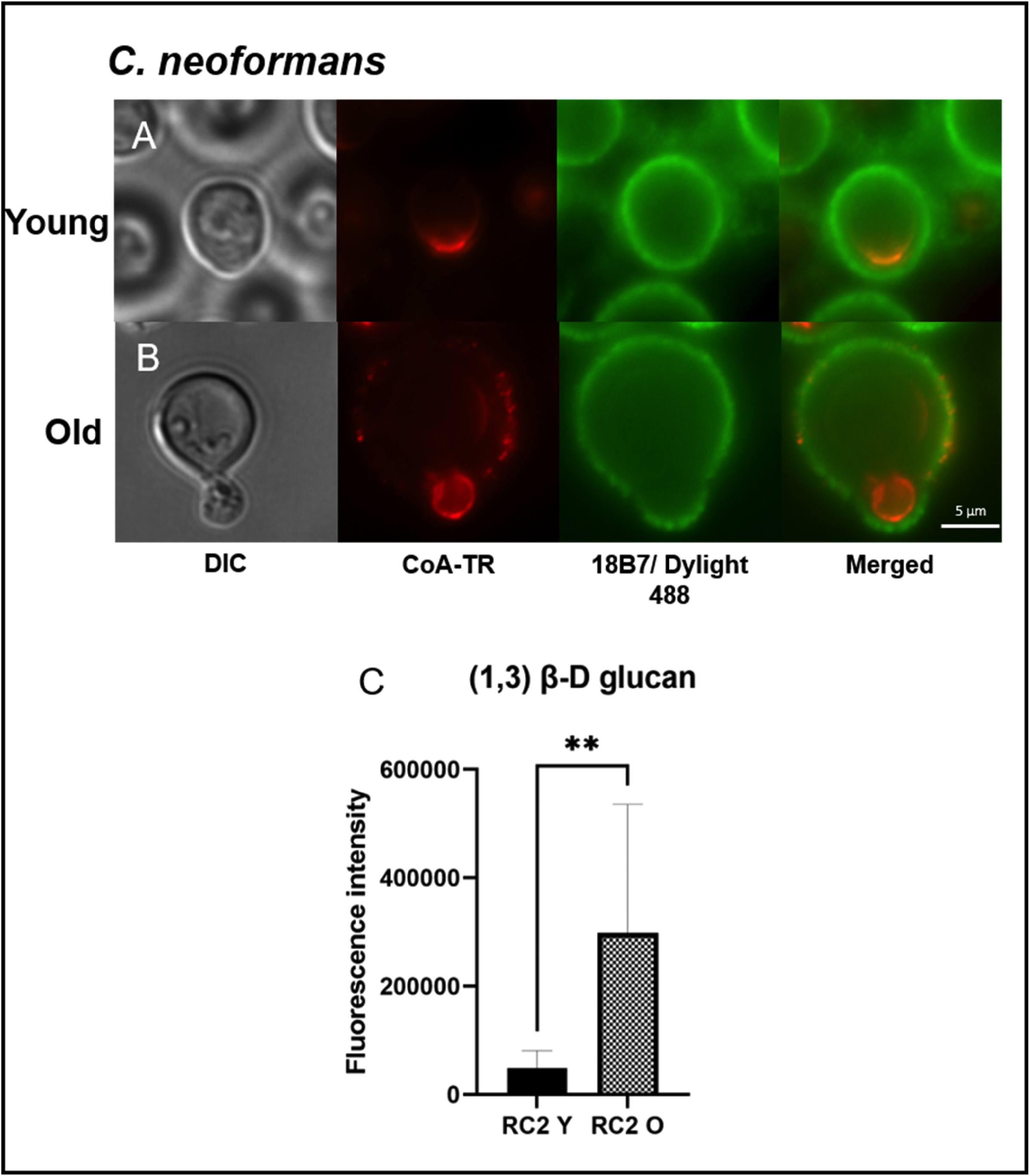
Co-localization of mannan in the polysaccharide capsule and enhanced content of b-glucan in the cell wall of old *C. neoformans* (*Cn*). Images represent microscopy analysis of RC2 *Cn* young (A) and old (B) cells. DIC (Differential Interference Contrast), followed by mannan (red) stained with concanavalin A conjugated to Texas red (CoA-TR), and GXM (green) in the capsule stained with primary monoclonal Ab 18B7 and secondary Ab anti-IgG conjugated to Dylight 488 (18B7/Dylight 488). Quantity of 1,3-β-D-glucan content of young cells (RC2 Y) and old *Cn* cells (RC2 O) were verified using biochemical assay with aniline blue (C). Experiments were done in biological triplicates, and statistical analysis were done by unpaired *t-*test (**, p= 0.0062).

### Old yeast cells presented increased levels of cell wall components

Flow cytometry was used to quantify the relative levels of cell wall carbohydrates in both young and old *Cn* and *Cg* (Fig.S1). Mean fluorescence intensity (MFI) levels confirmed increased chitin levels in old *Cn* (p <0.0001, Fig.4A) and *Cg* (p = 0.014, Fig.4B) compared to young cells. In agreement with microscopic findings, both older *Cn* and *Cg* exhibited significantly more chitooligomers quantity (p < 0.0001 and p = 0.0008, respectively, Fig.4C&D). Although chitooligomers can be quantified by labeling with WGA, CFW penetrates the cell surface more efficiently, and it also binds to other chitin derivates (28). As expected, chitosan levels were also significantly higher in old *Cn* when compared to young cells (p = 0.0058, Fig.4E).

**FIG 4.**
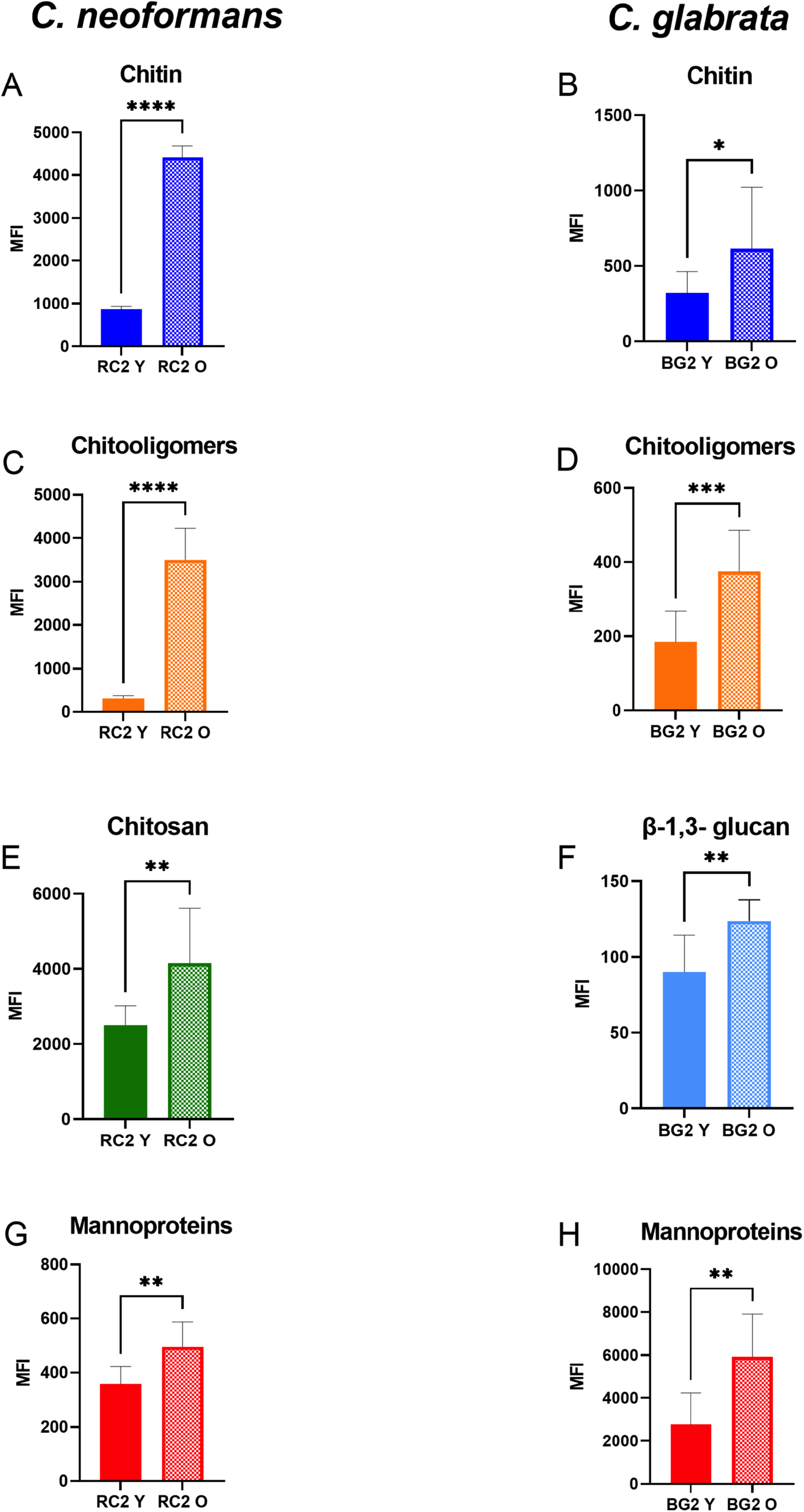
Old *C. neoformans* (*Cn*) and *C. glabrata* (*Cg*) presented increased cell wall content levels. By flow cytometry, analysis of levels in young (RC2 Y and BG2 Y) and old cells of *Cn* (RC2 O) and *Cg* (BG2 O) presented of total chitin (A and B), exposed chitooligomers (C and D), chitosan (E), β-1,3-glucan (F) and mannoproteins (G and H). Unstained cells were sorted as controls to determined positive labeling. For each group, a total of 10,000 events were gated and levels of chitin (blue; calcofluor white, CFW), chitooligomers (orange; WGA-TRITC), chitosan (green; Eosin Y, EY), β-glucan (light blue; Fc-Dectin1/Dylight 405), and mannan (red; concanavalin A, Texas Red conjugated) were represented by the mean fluorescence intensity (MFI). The following lasers were used: violet 405 nm (for CFW and Dylight 405), YelGreen 561 nm (for WGA-TRITC and CoA-TR), blue 488 nm (for EY). All experiments were done in biological triplicates and statistical analyses were performed by *t-*test (****, p ≤ 0.0001; *** p ≤ 0.001; ** p ≤ 0.01; * p ≤ 0.05).

Regarding the mannoproteins content, old *Cn* and *Cg* presented higher levels when compared to younger cells (p = 0.0019 and p = 0.0015, respectively, Fig.4G&H). β-D glucan content in *Cn* cell wall could only be quantified using a biochemical assay. These data indicated old *Cn* contained increased levels in comparison to young cells (p = 0.0062, Fig.3C). Also, higher level of β-glucan content was found in the cell wall of old *Cg* when compared to that in young cells (p= 0.0025, Fig.4F), which could not be documented by microscopy.

### Old yeast cells developed thicker and robust cell walls, giant vacuoles, and increased multivesicular bodies-like structures

As expected, TEM analysis showed that older *Cn* had a significantly thicker cell wall when compared to that of younger cells (235.9 vs. 123.7 nm; p < 0.0001). Specifically, we found in young *Cn*, the inner cell wall layer measured on average 66.76 nm, and the outer layer 91.29 nm. In old cells, the average thicknesses of the inner layer were 103.4 nm and grew to 138.3 nm (p < 0.0001), respectively. These results suggest that both cell wall layers equally enlarged during replicative aging. Old *Cg* also showed a notable growth in the total thickness of the cell wall when compared to young cells (132.5 vs. 93.11 nm; p < 0.0001). Here the inner cell wall layer was thicker than the outer, and both grew with age. Specifically, in old *Cg* the inner layer measured 103.5 nm vs. 78.21 nm in young cell (p = 0.0007). Conversely, the outer cell layer of young *Cg* was thicker than that of old *Cg* (34.75 nm vs. 25.39 nm; p = 0.0017) (Fig. 5B).

**FIG 5.**
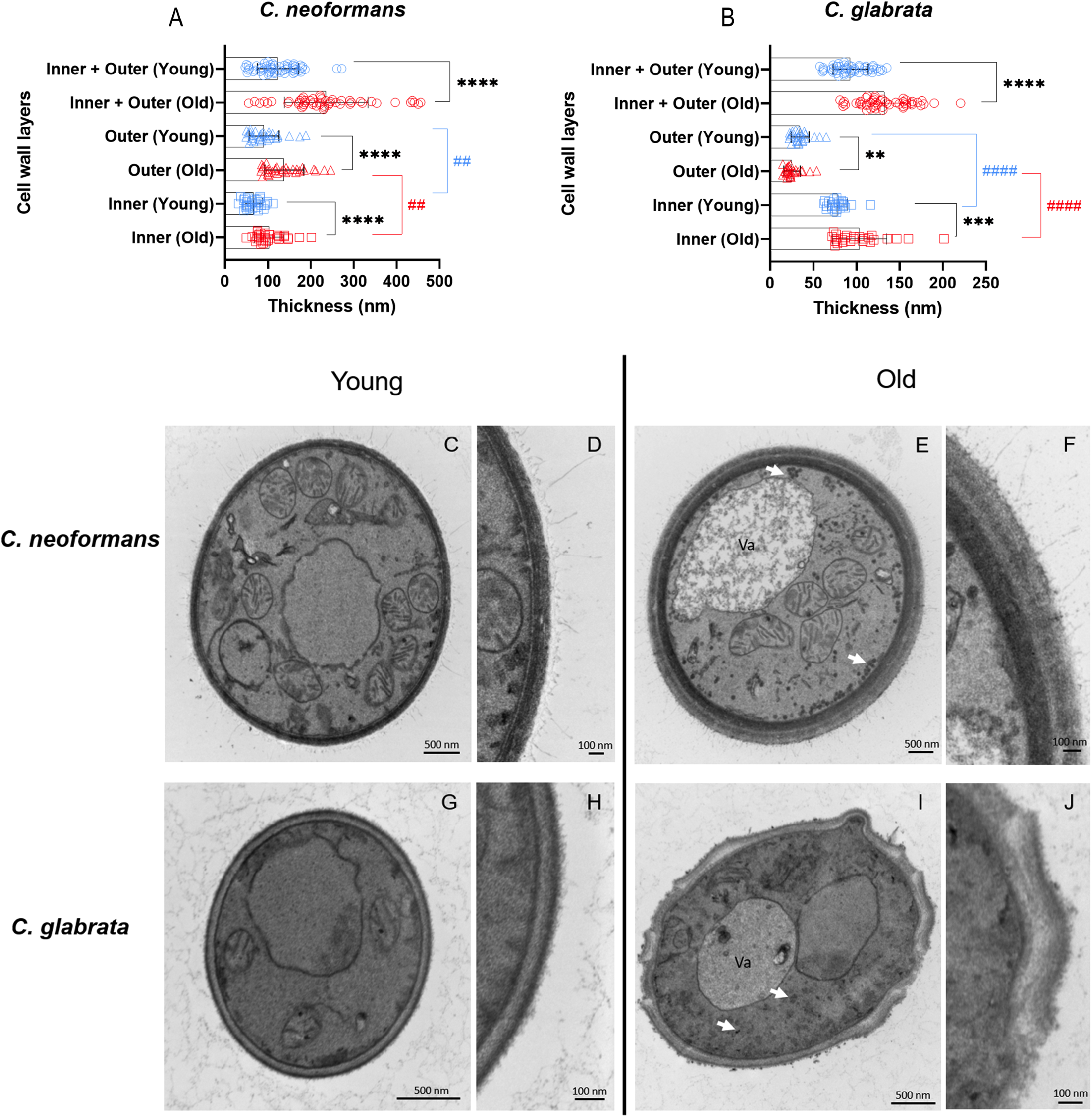
Old yeast cells present thicker cell walls and an accumulation of vesicle-like structures in the cytosol. Quantification of the thickness of the inner (squares), outer (triangles), and total (inner + outer) (circles) layers of young (blue) and old (red) *C. neoformans* (*Cn*) (A) and *C. glabrata* (*Cg*) (B). Data represent the analysis of 30 individual cells for each group by ImageJ/Fiji software. Unpaired *t*-test with Welch’s correction was used to compare the pairs (young and old) of each group, and error bars represent the standard deviation (****, p ≤ 0.0001; ***, p ≤ 0.001; **, p ≤ 0.01; ####, p ≤ 0.0001; ##, p ≤ 0.01). The asterisk symbol (*) represents differences between the groups young and old, whereas the pound symbol (#) represents differences within the groups young or old. Representative electron micrographs showing the ultrastructural changes of young and old cell walls of *Cn* (RC2) (C-F) and *Cg* (BG2) (G-J). White arrows point to vesicle-like particles in the cytoplasm of old yeast cells.

Next, we tested if replicative aging affected cell wall strength. Cell wall integrity is vital for maintaining cell wall functionality and structure (29). We assessed the sensitivity of old cells to cell wall stressors, including Congo red (inhibits assembly of cell wall polymers like chitin), caffeine (interferes with cell wall-related signal transduction pathways, and sodium dodecyl sulfate (disrupts plasma membrane) (28, 30). These data showed aging had no impact on resistance of the cell wall in old *Cn* and *Cg* (Fig.S2).

The ultramicroscopic images showed that old cells of both yeasts exhibited expanded vacuoles with amorphous lumen content, occupying a significant fraction of the total cell (Fig. 5E&I). Interestingly, the cytoplasm and cell wall (Fig.S3) of old cells contained a large number of electron-dense particles (Fig.5E&I), suggestive of vesicles derived from the conventional secretion pathway. The vacuoles in old *Cn* (Fig.6D) and *Cg* (Fig.6H) contained vesicles resembling multivesicular bodies (MVBs). MVB-like structures accumulated in 67.35% of old *Cn*, whereas they were found in only 32.69% of young cells. In a similar fashion in old *Cg*, MVB-like structures were present in 33.96% of old cells and only in 4% in young cells (Table 1).

**FIG 6.**
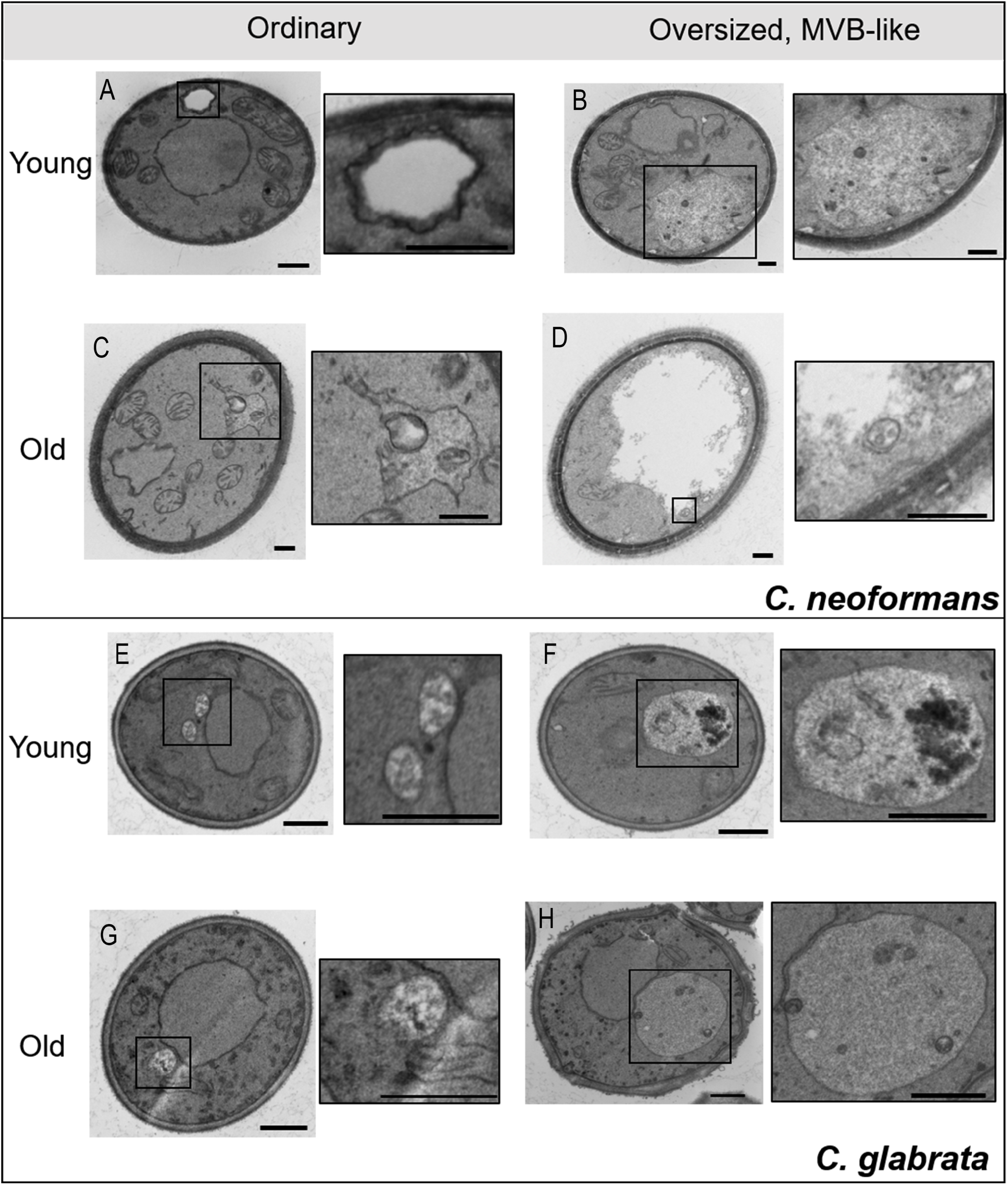
Alterations of vacuolar morphologies in young and old *C. neoformans* (*Cn*) and *C. glabrata* (*Cg*). Representative transmission electron microscopy (TEM) images of young and old cells of *Cn* in the upper panel (A-D), and *Cg* in the lower panel (E-H). Boxed areas illustrating vacuoles (ordinary or oversized, with or without intravacuolar vesicles) (left panels), were magnified in the right panels. Scale bars represent 500 nm.

**Table 1.** Distribution of multivesicular bodies (MVB)-like structures in young and old cells of C. neoformans (Cn) and C. glabrata (Cn).

| Strain | Vacuole | Young | Old |
| --- | --- | --- | --- |
| Cn (RC2) | Ordinary | 64.15% (34) | 32.69% (17) |
|  | Oversized, MVB-like | 24.53% (13) | 67.31% (35) |
|  | Other | 11.32% (6) | 0 |
| Cg (BG2) | Ordinary | 80% (40) | 49.05% (26) |
|  | Oversized, MVB-like | 4% (1) | 33.96% (18) |
|  | Other | 16% (8) | 16.99% (9) |

Transmission Electron Microscopy (TEM) images of at least 50 cells from each group were analyzed according to the size of the vacuole and the presence of MVB-like structures.

### Old *Cn* accumulated intracellular GXM and upregulated secretion gene *SEC14*

*Cn* extracellular vesicles (EVs) contain GXM (31). Analysis of ultrathin sections of both young and old *Cn* labeled with mAb18B7 for GXM (32) revealed a significant increase in GXM detection in the intracellular space in old compared to young *Cn* (Fig.7A-E). Immunogold labeled antibodies detected GXM in the vacuole (Fig.7D) and also bound to vesicle-like structures in the cell wall (Fig.7E) of old yeast. We also tested transcription of two genes essential for secretion *SEC14* and *SAV1* (33–35). *SEC14* transcription was upregulated (3.09 fold-change) whereas *SAV1* transcription was not significantly altered in old *Cn* (Table 2).

**FIG 7.**
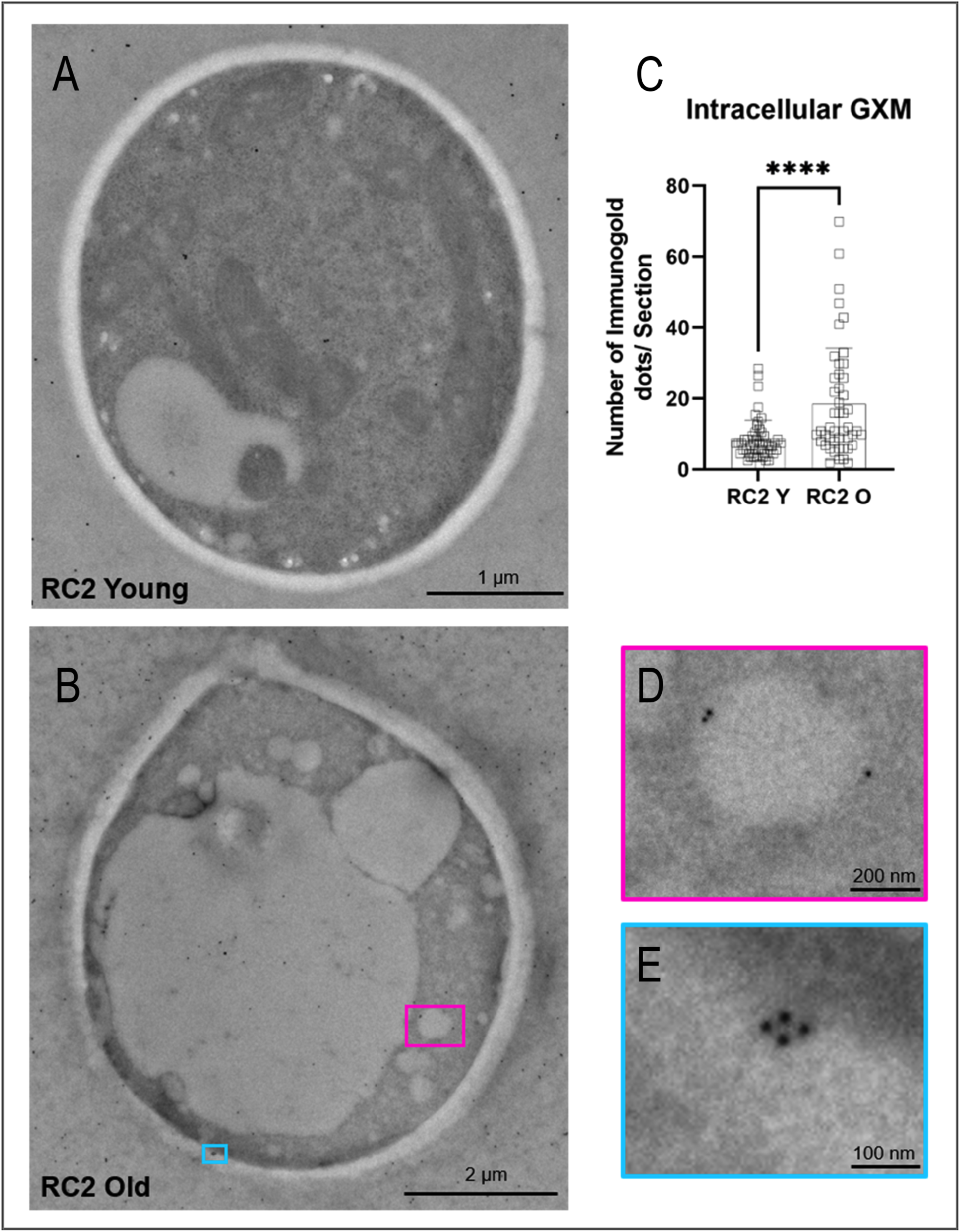
Intracellular GXM was enriched in old *C. neoformans* (*Cn*). Quantitative analysis of immunogold labelling to detect intracellular GXM with monoclonal antibody to GXM (mAb 18B7) in young (A) and old (B) *Cn* (RC2). In panel C, error bars represent standard deviations. Statistical significance was evaluated using unpaired *t*-test (****, p ≤ 0.0001). For each group (young and old), 50 cells were analyzed. A significantly increased amount of intracellular GXM was detected in old *Cn* compared to young cells. Illustrative images of old *Cn* cell showing the association of GXM inside vacuole (D) and with vesicular structures in the cell wall (E). Right panels in D (pink) and E (blue) represent magnifications of the boxed areas in left panel B.

**Table 2.**
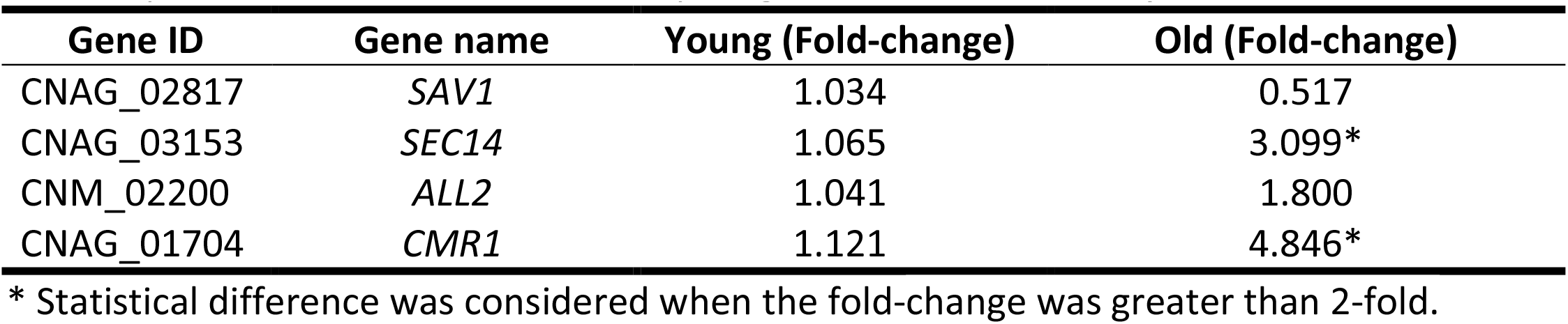
Expression of genes relevant for conventional secretion mechanisms, regulation of vacuole pH and calcium homeostasis in young and old cells of C. neoformans.

### The number, size, and pH of vacuoles are affected in yeast old cells

In fungi, the vacuole participates in secretory pathways, Ca^2+^ storage and pH regulation (36, 37). Because reduced vacuolar acidity has been reported in aged *S. cerevisiae* (38), we further investigated modifications in vacuole morphology and pH in the old cells. First, we stained the vacuolar membranes with FM™4-64 (Fig.8A&D). The average vacuole area was significantly larger in old compared to young cells, corresponding to an increase of 3.46-fold for *Cn* (0.7481 µm^2^ vs. 0.2162 µm^2^, p < 0.0001) and 4-fold in *Cg* (0.04 nm^2^ vs. 0.01 nm^2^, p < 0.0001). The surface area of old cells was 1.82-fold increased in *Cn* (41.38 µm^2^ vs. 22.68 µm^2^, p < 0.0001), and 1.63-fold in *Cg* (12.93 nm^2^ vs. 7.89 nm^2^, p < 0.0001) (Fig.S4).

**FIG 8.**
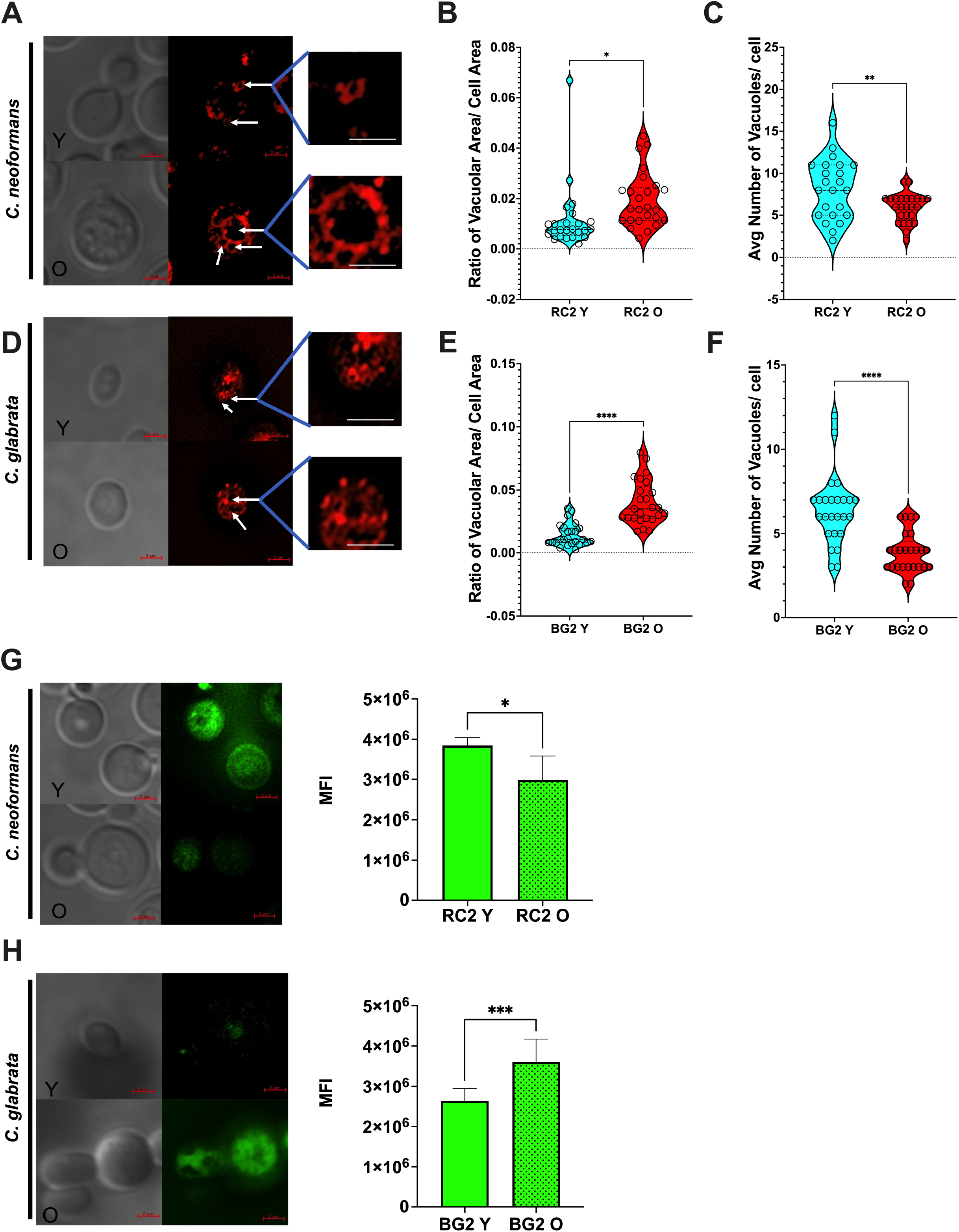
Vacuole morphology and pH are affected in yeast old cells. Images depicting vacuolar morphology (white arrows) stained with FM4-64 in young (Y) and old (O) cells of *C. neoformans* (*Cn*) (A) and *C. glabrata* (*Cg*) (D). DIC, differential interference contrast. The ratio vacuolar/cell area was significantly increased in old (red) *Cn* (B) and *Cg* (E) when compared to younger cells (blue). Unpaired *t*-test with Welch’s correction was used to generate the *p*-values (*, p = 0.0245; ****p < 0.0001). Average number of vacuoles per yeast cell was significantly lower in old (red) *Cn* (C) and *Cg* (F) when compared to respective young cells (blue). Unpaired *t*-test with Welch’s correction was used (**, p = 0.0098; ****, p < 0.0001). For all analysis, 25 cells of each group were measured using ImageJ/Fiji software. Measurement of quinacrine levels with a spectrophotometer showed a significantly decreased in fluorescence in aged *Cn* when compared to young cells (G). This was confirmed with microscopy. In contrast, significantly increased fluorescence was measured in old cells of *Cg* when compared to the young cells (H). This was also confirmed with microscopy. The assay was done in biological triplicate and unpaired *t*-test with Welch’s correction was used to generate the p-value (*, p = 0.0155; ***, p = 0.0007).

Calculations of the ratio of vacuole and cell size confirmed disproportional vacuolar growth in old *Cn* (p = 0.0155, Fig. 8B) and *Cg* (p < 0.0001, Fig.8E). The average number of vacuoles in young cells was higher in *Cn* (5.9 vs. 7.8 for old and young cells, p = 0.0231, Fig.8C) and *Cg* (3.9 vs. 6.4 for old and young, p < 0.001, Fig.8F). We hypothesize that the enlargement of vacuoles in aging cells could be the result of the fusion of multiple vacuoles.

Next, the vacuole pH of young and old cells was assessed with quinacrine staining, a fluorescent probe that increases under acidic conditions (39). In *Cn*, higher pH was observed in the vacuoles of old compared to young cells (3.85 x 10^6^ MFI vs 2.98 10^6^ MFI, p = 0.0155) (Fig.8G). Consistent with this finding *ALL2* expression, which is linked to the homeostasis of intracellular pH in *Cn* (40), trended up, and *CMR1,* which promotes calcium homeostasis (41), was significantly upregulated (4.84-fold increase, p < 0.005) (Table 2). In contrast, old *Cg* cells exhibited lower pH in vacuoles compared to young cells (3.6 x 10^6^ MFI vs 2.6 x 10^6^ MFI, p=0.0007) (Fig.8H).

A simplified illustration model with the phenotypic characteristics observed in this study during replicative aging in *Cn* and *Cg* are summarized in Fig. 9.

**FIG 9.**
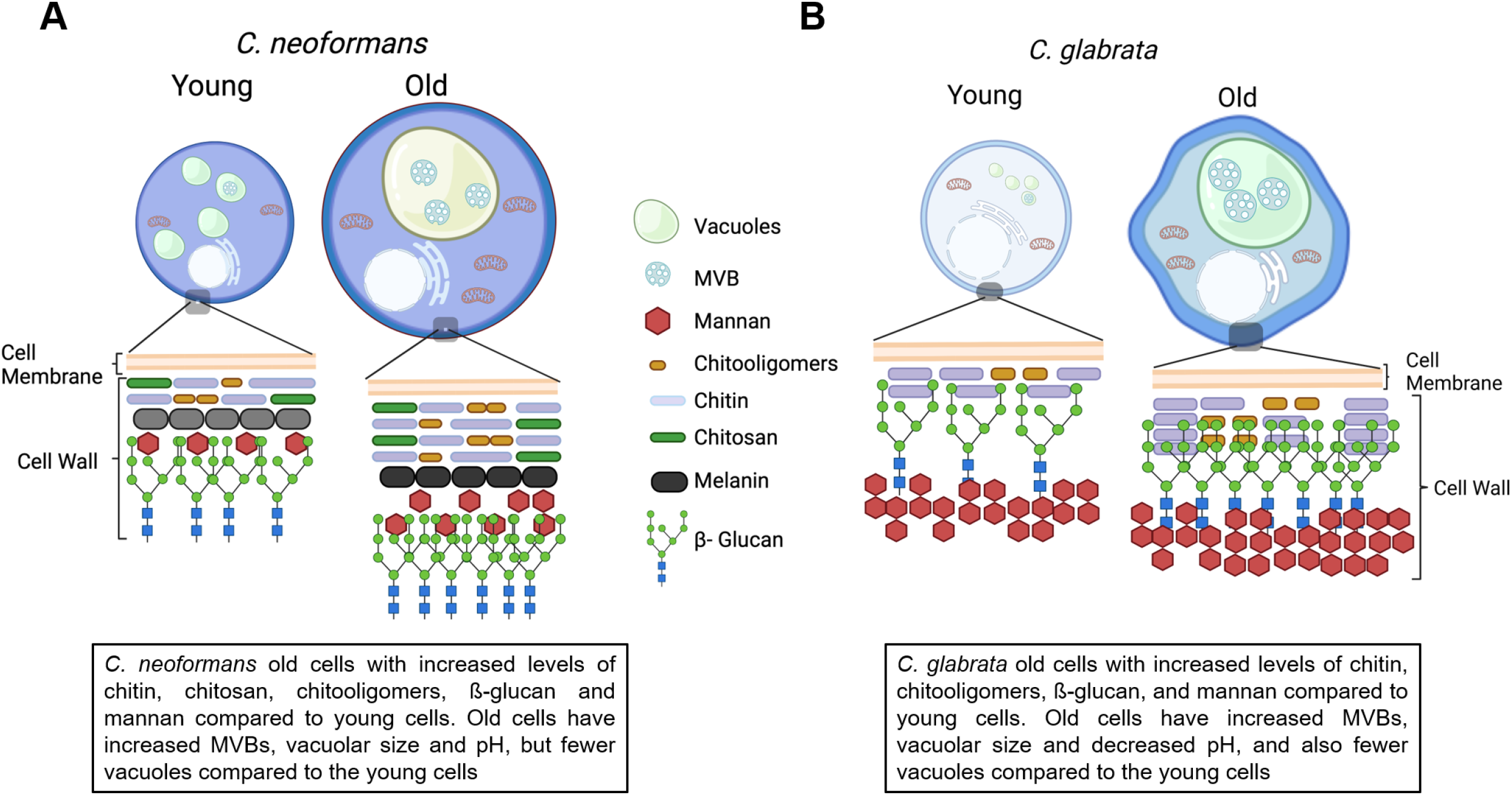
Illustrative summarization of phenotypic modifications observed during replicative aging in pathogenic yeast *C. neoformans* (A) and *C. glabrata* (B). During aging, cell wall increases in thickness, produce enlarged vacuoles and more multivesicular bodies (MVBs). Blue shapes indicate total chitin, dark green shapes indicate chitosan, orange shapes indicate chitooligomers, black shapes indicate melanin, red pentagon shapes indicate mannan, blue squares and green circles indicate β-glucan. The illustration was created using BioRender.

## DISCUSSION

The cell wall remains an attractive target for developing fungal vaccines (9) and antifungal drugs (10) because it is essential for fungal virulence (11). This study demonstrates that critical genes for cell wall synthesis are upregulated during replicative aging of *Cn* and *Cg*. The development of a thicker cell wall in old cells is probably facilitated by increased intracellular trafficking, associated with the formation of oversized vacuoles that contain MVB-like structures. Additionally, our results indicate that the morphology and pH of vacuoles are affected in old yeast cells. The observed reorganization of the cell wall partially explains the enhanced resistance to phagocytosis and antifungals, which has been observed in *Cn*, *Cg, C. albicans* and *C. auris* (21, 42–44).

Higher chitin levels result from budding and are implicated in increasing antifungal resistance, albeit not in all fungal species (45–47). For instance, chitin content is associated with caspofungin resistance in *Cn* (46), but not in *Cg* (45). Specifically, increasing chitin levels were noted throughout the lifespan in both yeasts similar to *S. cereviseae* (48). Staining patterns for chitooligomers mimicked the chitin staining pattern with more diffuse staining throughout the cell wall of old *Cn* and strictly bud scar-associated in *Cg*. In *Cn*, binding of chitooligomer-specific antibodies increases fungicidal efficacy of amphotericin B (49). As chitooligomers are fungi-specific, this novel therapeutic approach could be explored to treat the enhanced antifungal tolerance of old *Cn* and possibly *Cg*.

In *Cryptococcus* species, chitosan is a critical virulence factor and its levels can exceed chitin (16, 50, 51). It is produced from chitin by deacetylating enzymes (25), and *CDA3* transcription was upregulated in old *Cn*. Cda3 activity was more important during infection by *C. gattii* (50), however Cda1 (52) and Cda2 were the main regulators for *Cn* pathogenesis under host conditions (53). Chitosan inhibits protective Th1-type adaptive immune responses (54), maintains cell integrity, and allows melanization, thereby enhancing antifungal resistance (55–57). Our data suggests that higher chitosan content in the cell wall promotes enhanced melanization of old *Cn* cells during replicative aging (21). A common architecture for chitosan-containing elements in the fungal cell wall from ascomycetes to basidiomycetes has been proposed based on NMR studies (58). Although *Cg* produces a melanin-like pigment (59), it is unknown if it accumulates during replicative aging. Chitosan was not detected in any *Cg* cells similar to other *Candida* species, where expression is limited to chlamydospore (26).

Higher β-1,3-D-glucan levels were confirmed in old *Cn* by biochemical assays and in old *Cg* by fluorescent staining. Two of the three *FKS* genes involved in the synthesis were found to be upregulated in old *Cg* (42). However, *FKS1* levels in old *Cn* were not significantly increased. Transcript and protein levels do not always correlate, and unstable mRNA can be translated more efficiently (60). In *Cn* lacking post-transcriptional gene regulator *PUF4*, the *FKS1* mRNA and Fks1p levels did not correspond (46). Unfortunately, we were unable to quantify α-glucan with the available methods, but the α-glucan synthase *AGS* was upregulated in old *Cn*, indicating that this glucan is likely also increased. Nevertheless, it is still unclear whether the quantification of the cell wall content may be influenced by the enlargement of old cells. Further analysis, such as High-Performance Liquid Chromatography, would be required to resolve this issue (61, 62). However, the low yield of old cells isolated from yeast culture is still currently a limitation for performing this technique.

Fungal pathogens can evade host immune recognition by masking β-glucans at the cell wall surface (63, 64). The exposure of β-glucans is reduced by *Cn’s* capsule (61), and by mannan in the outer cell wall layer in *Candida* species (45, 65). Mannan is present in *Cn* polysaccharide capsule (66, 67), thus we propose that the observed accumulation of mannan could be due to augmented vesicle export. Alternatively, an altered permeability in the polysaccharide capsule in old cells (68) may efficiently expose the mannan epitopes. Altered mannoprotein content may contribute to the poor recognition of old yeast by host phagocytes (21, 43), which is suggested by data from *C. gattii* (69) and *C. albicans* (70).

Comparisons of TEM-images of *Cn* and *Cg* identified differential growth of their inner and outer cell wall layer. The outer cell wall layer was thicker in old *Cn*, although the inner cell wall layer dominated in *Cg*. Chitinous structures and chitosan are evenly distributed across the cell wall of *Cn* but must be flexible enough to permit even expansion of the cell wall. We hypothesize the heterogeneous composition of the inner cell wall in *Cg* result from chitin breaks in the bud scars, similar to *S. cerevisiae* (48) and causes the wavy cell wall. It is conceivable that the mannan layer in old *Cg* is more compactly arranged than in young cells. Reduced fibril length on the cell surface has been correlated with increased exposure on β- 1,3-glucan (45). Alternatively, the outer layer of aged *Cg* can be more fragile and could have partially detached during ultrastructural processing. The increase in the inner cell wall layer thickness avoids a hyperosmotic shock and favors fungal cell survival (71). Most importantly, analogous to *S. cerevisiae* (48), old and young cells exhibit comparable tolerance to common cell wall stressors. Furthermore, comparable doubling times of 10GEN old *Cn* as well as 14GEN old *Cg* cells (43, 72) indicate that old cells are equally fit in the host evironment.

Another important discovery is the development of oversized vacuoles with multiple vesicle-like structures in the cytoplasm of old yeast cells. Enlarged vacuoles have been described in old as well as nutrient-deprived *S. cerevisiae* cells (37, 38, 73). Vacuoles play a role in membrane trafficking (74) and protein sorting (37). Secretion mechanisms can also be influenced by autophagic processes (75). The autophagosome can merge first with MVBs and then with the cell membrane to deliver heterogeneous cargo outside the cell (76). In *Cn*, MVBs participate in laccase trafficking (77), a cell-wall associated virulence factor(78), as well as GXM and melanin are exported through EVs (31, 79). Cytoplasmic MVBs can originate fungal EVs (80). Cell wall architecture proteins were found in EVs from *Cn* (81), *Cg* (82), *S. cerevisiae* (83), *Histoplasma capsulatum* (84), and *Aspergillus fumigatus* (85). The cloning of a “mother cell enriching mutant” in *Cn* and *Cg*, similar to the one developed in *S. cerevisiae* (86), would permit prolonged growth of old cells, with isolation and analysis of their EV content.

The enlarged vacuole in old *Cn* and *Cg* cells is most likely the result of the fusion of smaller vacuoles. Vacuolar fusion in *S. cerevisiae* is necessary and sufficient for lifespan extension (87). Vacuolar acidification is also critical determinant of replicative lifespan and declines continuously with age, thereby limiting the replicative lifespan of a cell (88, 89). In *C. albicans*, vacuolar pH is a regulator of fission and fusion of vacuoles (90). We demonstrated loss of vacuolar acidity in old compared to young *Cn* similar to *S. cerevisiae* (38) but different from old *Cg*, where the vacuole pH was more acidic. Different factors including glucose starvation can also affect vacuolar pH (91). It is possible that *Cg* is experiencing more carbon starvation and has not advanced in its life span as much as *Cn*. Differences in pH regulation during aging and carbon starvation have been described in different *Cn* strains (92). *Cn* has a unique protein (All2p) involved in pH homeostasis (93), which accumulates during aging in low glucose conditions (40). More extensive analysis will be necessary to better understand the mechanisms involved in age-associated regulation of intracellular pH in these yeast cells, including the use of other pH probes that consistently co-localize within the vacuoles (94). It is noteworthy that these studies were conducted on a single strain for each pathogen and, therefore, further work including several strains of different genotypes is required.

In summary, these data provide insights regarding the dynamic of cell wall rearrangement during aging of *Cn* and *Cg*. The data highlight the divergence between these pathogens in budscar formation throughout the life span and uncovers differences such as cell wall composition, especially concerning mannan and chitosan. This work opens new avenues of investigation, such as the impact of cell wall reorganization in old yeast cells on the host innate response and sensitivity to cell wall targeting antifungals. Further work is also needed to elucidate the implications of EVs secretion during aging, including cargo and “cross-talk” between young and old pathogenic yeast cells and explore the association among aging, intracellular pH, and mitochondrial function.

## MATERIAL AND METHODS

### Strains and media

Fungal strains *Cn* RC2 (ATCC 24067 variant) and *Cg* BG2 were kept on YPD agar (Difco BD) plates. Synthetic media (SM) (44) was used for cultivation of yeast cells.

### Isolation of young and old *Cn* and *Cg* cells

Yeast cells (10^8^) from an overnight culture were washed with PBS and labeled with 8mg/ml sulfosuccinimidyl-6-[biotin-amido] hexanoate (Sulfo-NHS-LC-LC-Biotin, 21338, ThermoFisher Scientific) for 30 minutes at room temperature (RT). Subsequently, cells were washed with PBS and grown 5-7 doubling times. Then, 100 µl of magnetic streptavidin microbeads (130-048-101, Miltenyi Biotec) were added to every 10^8^ cells/mL, following incubation for 15 minutes at 4°C. Biotin-labeled yeasts were isolated using autoMACS magnetic columns (130-021-101, Miltenyi Biotec) and the autoMACS® Pro Separator (Miltenyi Biotec). The biotin-streptavidin labeled older cells were passed through a magnet where they get stuck while the younger unlabeled population flows through. The labeled cells are then recovered after the magnetic field is removed. The positive cell fraction was grown again in SM media to the desired generation (10GEN *Cn*, 14GEN *Cg*) and isolated as outlined above. Young cells washed off from the magnetic columns were kept as controls. The purity of fractions was verified by microscopy. In this work, “old” *Cn* are 10-generations old, and “old” *Cg* are 14-generations old.

### RNA isolation and quantitative reverse transcriptase polymerase chain reaction (RT-qPCR)

RNA isolation and RT-qPCR was done as described (22). Briefly, the RNAeasy kit (Qiagen) was used with on-column DNAse digestion, following the manufacturer’s instructions. The concentration of total RNA was measured using nanodrop (Eppendorf). The quantity of RNA was adjusted to 250 ng and converted to cDNA using Verso cDNA synthesis kit (Thermo Scientific). cDNA was diluted and used in a mix that contained Power SYBR® Green Mastermix (Applied Biosystems), together with oligo(dT) primers (Table S1).

The experiment was performed using the High-Performance Real Time PCR (Roche) LightCycler® 480 Systems (Roche). Gene expression was computed using ΔΔCt, *ACT1* was used to normalize the gene expression, and the gene expression in young cells were used as the reference. Values above two-fold were significant. C_t_ values of *ACT1* in old and young cells were similar to one another, signifying that *ACT1* gene expression does not vary between the young and old cells. Each experiment was repeated at least twice in two separate days.

### Cell wall staining

Young and old cells (10^6^) were fixed in 4% paraformaldehyde (PFA) for 15 min at RT, washed with PBS, and stained with calcofluor white (CFW; cell wall staining of chitin; 5 μg/ml for 10 minutes at 37°C), Concanavalin A, Texas Red conjugate (ConA-Texas red; cell wall staining of mannoproteins; 50 μg/ml for 45 minutes at 37°C), Eosin Y (EY, cell wall staining of chitosan; 300 μg/ml for 10 minutes at 37°C) (95) or tetramethylrhodamine-labeled wheat germ agglutinin (TRITC-WGA; cell wall staining of chitooligomers; 5 μg/ml for 30 minutes at 37°C) (14).

To stain β−glucan and GXM, fungal cells were blocked with 1% bovine serum albumin (BSA) for 1h at 37°C and incubated respectively with the primary antibody Fc-Dectin 1 (AdipoGen AG-40B-0138) (5 μg/ml; 40 minutes at 4°C) (28) or mAb 18B7 (10μg/mL; 1h at 37°C) (32). After washing with PBS, cells were incubated with secondary antibody DyLight 405 goat anti-human IgG + IgM (5 μg/ml; 30 minutes at 4°C), or with an anti-murine IgG Dylight 488 (10μg/mL; 30 minutes at 37°C) (Jackson ImmunoResearch).

Prior to analysis using a BD LRSForthessa™ flow cytometer, cells were washed and resuspended in PBS. The following lasers were used: blue, 488 nm (for: EY); YelGreen, 561 nm (for: ConA-Texas red and WGA-TRITC); violet, 405 nm (for: CFW and 405 Dylight). Data were analyzed using BD FACSDiva™ or FlowJo v10.1 software (FlowJo, LLC). A total of 10,000 events were gated in the forward scatter/ side scatter (FSC/SSC) plots and represented as histograms with mean fluorescence intensity (MFI) on the x-axis and cell counts on the y-axis. Unstained cells were used as negative controls. For fluorescence microscopy, the respective channels were used: FITC (EY and Dylight 488), DAPI (CFW and Dylight 405) and DsRed (Texas red and TRITC). Imaging was performed at 100x in a Zeiss Axio Observer or a Zeiss Axiovert 200M (Thornwood, NY). The same exposure time was used to image young and old cells. The images were processed using ImageJ software.

### (1,3) β-D glucan quantification

*Cn* (4 x 10^6^) were washed with Tris-EDTA buffer (pH 8.0) and suspended in 500 µl of Tris-EDTA and 56 µl NaOH (10 M), to achieve a final concentration of 1M. To solubilize (1,3) β-D glucan, cells were incubated at 80°C for 30 minutes. After, 2.1 ml of aniline blue solution (0.03% aniline blue, 0.18M HCl, 0.49M glycine/NaOH pH 9.5) was added to the cells to interact with linear (1,3) β-D glucan. Following homogenization, cells were incubated for 30 minutes at 50°C and for 30 minutes at RT. The fluorescence was measured using a plate reader SpectraMax i3X (Molecular Probes), excitation at 400 nm and emission at 460 mn (96).

### Ultramicroscopy

Cells were fixed in 3% glutaraldehyde in 0.1M sodium cacodylate buffer pH 7.4 at 4°C overnight. Then, cells were washed 3x with 0.1M sodium cacodylate buffer and dispersed in ultra-low temperature agarose at 4°C for 30 min. Subsequently, the agarose samples were cut into small cubes, and the pieces were post-fixed in aqueous 1% KMNO_4_ for 1h at RT. Blocks were rinsed with dH_2_O and treated with 0.5% sodium meta-periodate for 15 minutes at RT to allow infiltration. Next, blocks were rinsed with dH_2_O and dehydrated through a graded ethanol series. Samples were treated with 100% propylene oxide, embedded with Spurr Resin and polymerized in an oven at 60°C for 24h. Ultrathin sections (80 nm) were cut using a Leica EM UC7 Ultramicrotome, placed on 300 mesh copper grids, and counterstained with uranyl acetate and lead citrate. Images were acquired at magnifications x 18,500 and x 49,000 using an FEI TeCnai 12 BioTwinG^2^ TEM. Digital images were acquired with an AMT XR 60 CCD digital camera system. At least 30 young and old cells were analyzed, and the thickness of cell wall layers were measured by ImageJ software.

### Intracellular glucuronoxylomannan (GXM) of *Cn*

*Cn* cells were fixed for 45 minutes at RT in Fixate solution: Synthetic media + 4% PFA + 0.1% glutaraldehyde + 0.1M Cacodylate buffer. Samples were washed twice with 0.1M Cacodylate buffer, suspended in 0.1M Cacodylate buffer and left overnight at 4°C. The cells were washed with PBS, placed in gelatin, cut into cubes of 1 mm, and postfixed in 0.1 M sodium cacodylate containing 2.0% paraformaldehyde. Samples were then dehydrated through a graded series of ethanol, with a progressive lowering of the temperature to -50°C in a Leica EMAFS, embedded in Lowicryl HM-20 mono-step resin (Electron Microscopy Sciences), and polymerized using UV light. Ultrathin sections were cut on a Leica EMUC7, immunolabeled with mAb 18B7 (50 µg/ml) and gold-labelled anti-mouse IgG. Control systems included ultrathin sections incubated without the mAb 18B7, followed by the gold-labeled anti-mouse IgG. Samples were stained with uranyl acetate and viewed on a JEOL 1400 Plus transmission electron microscope at 80kv. Gold particles in electron micrographs were quantified as described (97). For quantitative analysis of GXM labeling, 50 cells of the young group and 47 cells from the old group were analyzed. For normalization, the average number of gold particles counted in control systems (no mAb 18B7 labeling) was subtracted from the total number of gold particles of each mAb 18B7-labeled section, as described (98).

### Vacuole morphology and pH analysis

Vacuoles from 10^6^ fungal cells were stained at 37°C with FM™4-64 (ThermoFisher) (10 µg/ml) for 10 minutes, following manufacturer’s protocol. Another set of 10^6^ fungal cells were labeled with quinacrine in separate tubes at a final concentration of 200 μM and incubated for 20 mins at 37°C. Cells were washed three times with PBS and imaged at 100x using a deconvolution microscope (Zeiss Axiovert 200M), using the filters Texas Red (for FM4-64) and GFP (for quinacrine). Twenty-five cells from each group and strain were analyzed using ImageJ software.

Also, 200 µl of yeast suspensions (10^6^) labeled with quinacrine were plated in triplicates in a black 96-well plate, and the fluorescence was measured with a spectrophotometer Spectramax i3X (Molecular Probes), using excitation at 436 nm and emission at 525 nm. The values of unstained cells were subtracted from the values obtained from the stained cells. All experiments were repeated three times independently.

### Cell wall stressors assay

Serial dilutions of yeast suspensions (final concentrations 10^5^ to 10^2^ cells) were spotted on YPD agar containing each of the following stressors: fluorescent brightener 28 (FB 28) 1.5 mg/ml; Congo Red 0.5%, caffeine 1 mg/ml or 0.5 mg/ml, Sodium Dodecyl Sulfate (SDS) 0.01% or 0.0025%. Next, the plates were incubated at 30°C and 37°C for 48h. Control conditions included cells added on YPD agar without the stressors. For Congo Red, the stressor was added to the YPD medium prior to autoclaving. Whereas for caffeine, FB 28 and SDS, stock solutions were prepared, filter sterilized and added to YPD after autoclaving (28).

### Statistical analysis

Statistical differences between young and old cells were performed by *t*-test with Welch’s correction. All statistical tests we performed with GraphPad Prism 9.0 software (GraphPad Software, Inc.).

## ACKNOWLEDGMENTS

We would like to acknowledge TEM sample preparation and imaging was performed at the *C-MIC: Central Microscopy Imaging Center* at Stony Brook University, Stony Brook, New York, and alternatively at the Analytical Imaging Facility, Albert Einstein College of Medicine, Bronx, New York. We also are thankful to the Flow Cytometry Research Core Facility at Stony Brook Hospital and Dr. Del Poeta for allowing us to use the fluorescent microscope. mAb 18b7 was a gift from Dr. Arturo Casadevall.

This work was financially supported by the National Institutes of Health (NIH 1R01-AI127704-01A1 to B.C.F). The contents of this article do not represent the views of VA or the United States Government.

---

**Table S1.**
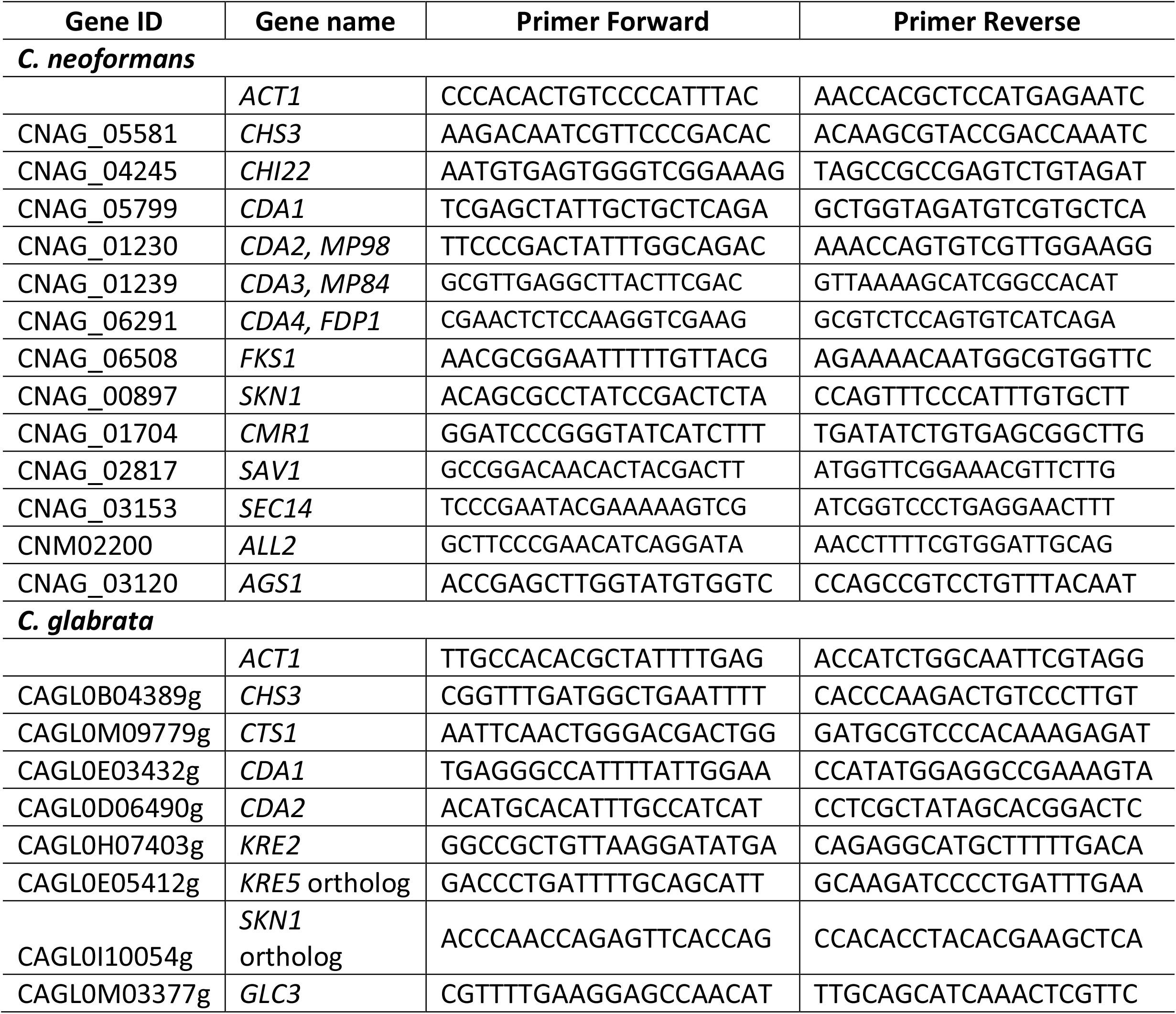
Sequence of primers (5’-3’) used in this study.

**FIG S1.** Flow cytometry analysis of young and old *C. neoformans* and *C. glabrata c*ells. At least 10,000 events were gated in the FSC/SSC plots and are represented as histograms with mean fluorescence on the x-axis and cell counts on the y-axis. Unstained young (red) and old (green) cells were sorted as controls to determine positive labeling of young (blue) and old (orange) cells. CFW (Calcofluor White, chitin), WGA-TRITC (tetramethylrhodamine-labeled wheat germ agglutinin, chitooligomers), EY (Eosin Y, chitosan), CoA-Texas Red (Concanavalin A conjugated to Texas Red, Mannan), Fc-Dectin1/Dylight 405 (β-glucan). Data were analyzed by FlowJo V10 Software and the image is representative of one experiment producing similar results that were obtained in three independent experiments.

**FIG S2.** No impact on cell wall integrity was observed in old *C. neoformans* (*Cn*) and *C. glabrata* (*Cg*) cells. Young and old cells of *Cn* and *Cg* were serially diluted and spotted onto YPD with the addition of several cell wall stressors (caffeine; FB 28, fluorescent brightener 28; congo red and SDS, sodium dodecyl sulfate) at 30°C and 37°C for 48h. This experiment was repeated twice with similar results.

**FIG S3.** Ultramicroscopic image of old *C. neoformans* cells with numerous vesicle-like particles in the cytosol, inside the vacuole and crossing the cell wall indicated by white arrows. Scale bar represents 500 nm.

**FIG S4.** Cell surface and vacuolar areas are significantly increased in old *C. glabrata* (A and B) and *C. neoformans* (C and D). Fungal cells were stained with FM4-64 and analyses of 25 cells for each group (young and old) and strain were measured using ImageJ/Fiji software. Statistical analyses were performed by *t*-test with Welch’s correction (****, p ≤ 0.0001).

## References

1. Vallabhaneni S, Mody RK, Walker T, Chiller T. 2016. The Global Burden of Fungal Diseases. Infect Dis Clin North Am 30:1–11.

2. Nnadi NE, Carter DA. 2021. Climate change and the emergence of fungal pathogens. PLOS Pathogens 17:e1009503.

3. Casadevall A. 2020. Climate change brings the specter of new infectious diseases. The Journal of Clinical Investigation 130:553–555.

4. Bongomin F, Gago S, Oladele RO, Denning DW. 2017. Global and Multi-National Prevalence of Fungal Diseases-Estimate Precision. Journal of fungi (Basel, Switzerland) 3:57.

5. Rajasingham R, Smith RM, Park BJ, Jarvis JN, Govender NP, Chiller TM, Denning DW, Loyse A, Boulware DR. 2017. Global burden of disease of HIV-associated cryptococcal meningitis: an updated analysis. Lancet Infect Dis 17:873–881.

6. Healey KR, Perlin DS. 2018. Fungal Resistance to Echinocandins and the MDR Phenomenon in Candida glabrata. J Fungi (Basel) 4.

7. Pappas PG, Lionakis MS, Arendrup MC, Ostrosky-Zeichner L, Kullberg BJ. 2018. Invasive candidiasis. Nature Reviews Disease Primers 4:18026.

8. Perfect JR. 2017. The antifungal pipeline: a reality check. Nature reviews Drug discovery 16:603–616.

9. Oliveira LVN, Wang R, Specht CA, Levitz SM. 2021. Vaccines for human fungal diseases: close but still a long way to go. npj Vaccines 6:33.

10. Ibe C, Munro CA. 2021. Fungal cell wall: An underexploited target for antifungal therapies. PLOS Pathogens 17:e1009470.

11. Garcia-Rubio R, de Oliveira HC, Rivera J, Trevijano-Contador N. 2020. The Fungal Cell Wall: Candida, Cryptococcus, and Aspergillus Species. Frontiers in Microbiology 10.

12. Gow NAR, Latge JP, Munro CA. 2017. The Fungal Cell Wall: Structure, Biosynthesis, and Function. Microbiol Spectr 5.

13. Ruiz-Herrera J, Ortiz-Castellanos L. 2019. Cell wall glucans of fungi. A review. Cell Surf 5:100022.

14. Rodrigues ML, Alvarez M, Fonseca FL, Casadevall A. 2008. Binding of the wheat germ lectin to Cryptococcus neoformans suggests an association of chitinlike structures with yeast budding and capsular glucuronoxylomannan. Eukaryot Cell 7:602–9.

15. Fonseca FL, Guimarães AJ, Kmetzsch L, Dutra FF, Silva FD, Taborda CP, Araujo Gde S, Frases S, Staats CC, Bozza MT, Schrank A, Vainstein MH, Nimrichter L, Casadevall A, Rodrigues ML. 2013. Binding of the wheat germ lectin to Cryptococcus neoformans chitooligomers affects multiple mechanisms required for fungal pathogenesis. Fungal Genet Biol 60:64–73.

16. Banks IR, Specht CA, Donlin MJ, Gerik KJ, Levitz SM, Lodge JK. 2005. A chitin synthase and its regulator protein are critical for chitosan production and growth of the fungal pathogen Cryptococcus neoformans. Eukaryot Cell 4:1902–12.

17. Hembach L, Bonin M, Gorzelanny C, Moerschbacher BM. 2020. Unique subsite specificity and potential natural function of a chitosan deacetylase from the human pathogen Cryptococcus neoformans. Proc Natl Acad Sci U S A 117:3551–3559.

18. Free SJ. 2013. Fungal cell wall organization and biosynthesis. Adv Genet 81:33–82.

19. Sherrington SL, Sorsby E, Mahtey N, Kumwenda P, Lenardon MD, Brown I, Ballou ER, MacCallum DM, Hall RA. 2017. Adaptation of Candida albicans to environmental pH induces cell wall remodelling and enhances innate immune recognition. PLoS pathogens 13:e1006403–e1006403.

20. Bhattacharya S, Bouklas T, Fries BC. 2020. Replicative Aging in Pathogenic Fungi. J Fungi (Basel) 7.

21. Orner EP, Bhattacharya S, Kalenja K, Hayden D, Del Poeta M, Fries BC. 2019. Cell Wall-Associated Virulence Factors Contribute to Increased Resilience of Old Cryptococcus neoformans Cells. Front Microbiol 10:2513.

22. Bouklas T, Alonso-Crisóstomo L, Székely T, Jr., Diago-Navarro E, Orner EP, Smith K, Munshi MA, Del Poeta M, Balázsi G, Fries BC. 2017. Generational distribution of a Candida glabrata population: Resilient old cells prevail, while younger cells dominate in the vulnerable host. PLoS pathogens 13:e1006355–e1006355.

23. Charlet R, Pruvost Y, Tumba G, Istel F, Poulain D, Kuchler K, Sendid B, Jawhara S. 2018. Remodeling of the Candida glabrata cell wall in the gastrointestinal tract affects the gut microbiota and the immune response. Scientific Reports 8:3316.

24. Baker LG, Specht CA, Lodge JK. 2009. Chitinases are essential for sexual development but not vegetative growth in Cryptococcus neoformans. Eukaryot Cell 8:1692–705.

25. Baker LG, Specht CA, Donlin MJ, Lodge JK. 2007. Chitosan, the deacetylated form of chitin, is necessary for cell wall integrity in Cryptococcus neoformans. Eukaryot Cell 6:855–67.

26. Bemena LD, Min K, Konopka JB, Neiman AM. 2021. A Conserved Machinery Underlies the Synthesis of a Chitosan Layer in the Candida Chlamydospore Cell Wall. mSphere 6.

27. Reese AJ, Doering TL. 2003. Cell wall alpha-1,3-glucan is required to anchor the Cryptococcus neoformans capsule. Mol Microbiol 50:1401–9.

28. Esher SK, Ost KS, Kohlbrenner MA, Pianalto KM, Telzrow CL, Campuzano A, Nichols CB, Munro C, Wormley FL, Jr., Alspaugh JA. 2018. Defects in intracellular trafficking of fungal cell wall synthases lead to aberrant host immune recognition. PLOS Pathogens 14:e1007126.

29. de Oliveira HC, Rossi SA, García-Barbazán I, Zaragoza Ó, Trevijano-Contador N. 2021. Cell Wall Integrity Pathway Involved in Morphogenesis, Virulence and Antifungal Susceptibility in Cryptococcus neoformans. Journal of Fungi 7:831.

30. Ram AFJ, Klis FM. 2006. Identification of fungal cell wall mutants using susceptibility assays based on Calcofluor white and Congo red. Nature Protocols 1:2253–2256.

31. Rodrigues ML, Nimrichter L, Oliveira DL, Frases S, Miranda K, Zaragoza O, Alvarez M, Nakouzi A, Feldmesser M, Casadevall A. 2007. Vesicular polysaccharide export in Cryptococcus neoformans is a eukaryotic solution to the problem of fungal trans-cell wall transport. Eukaryot Cell 6:48–59.

32. Casadevall A, Cleare W, Feldmesser M, Glatman-Freedman A, Goldman DL, Kozel TR, Lendvai N, Mukherjee J, Pirofski LA, Rivera J, Rosas AL, Scharff MD, Valadon P, Westin K, Zhong Z. 1998. Characterization of a murine monoclonal antibody to Cryptococcus neoformans polysaccharide that is a candidate for human therapeutic studies. Antimicrobial agents and chemotherapy 42:1437–1446.

33. Yoneda A, Doering TL. 2009. An unusual organelle in Cryptococcus neoformans links luminal pH and capsule biosynthesis. Fungal Genet Biol 46:682–7.

34. Chayakulkeeree M, Johnston SA, Oei JB, Lev S, Williamson PR, Wilson CF, Zuo X, Leal AL, Vainstein MH, Meyer W, Sorrell TC, May RC, Djordjevic JT. 2011. SEC14 is a specific requirement for secretion of phospholipase B1 and pathogenicity of Cryptococcus neoformans. Mol Microbiol 80:1088–101.

35. Yoneda A, Doering TL. 2006. A eukaryotic capsular polysaccharide is synthesized intracellularly and secreted via exocytosis. Molecular biology of the cell 17:5131–5140.

36. Klionsky DJ, Herman PK, Emr SD. 1990. The fungal vacuole: composition, function, and biogenesis. Microbiol Rev 54:266–92.

37. Veses V, Richards A, Gow NAR. 2008. Vacuoles and fungal biology. Current Opinion in Microbiology 11:503–510.

38. Henderson KA, Hughes AL, Gottschling DE. 2014. Mother-daughter asymmetry of pH underlies aging and rejuvenation in yeast. eLife 3:e03504.

39. Pierzyńska-Mach A, Janowski PA, Dobrucki JW. 2014. Evaluation of acridine orange, LysoTracker Red, and quinacrine as fluorescent probes for long-term tracking of acidic vesicles. Cytometry A 85:729–37.

40. Orner EP, Zhang P, Jo MC, Bhattacharya S, Qin L, Fries BC. 2019. High-Throughput Yeast Aging Analysis for Cryptococcus (HYAAC) microfluidic device streamlines aging studies in Cryptococcus neoformans. Communications Biology 2:256.

41. Cao C, Wang Y, Husain S, Soteropoulos P, Xue C. 2019. A Mechanosensitive Channel Governs Lipid Flippase-Mediated Echinocandin Resistance in Cryptococcus neoformans. mBio 10.

42. Bhattacharya S, Fries BC. 2018. Enhanced Efflux Pump Activity in Old *Candida glabrata* Cells. Antimicrobial Agents and Chemotherapy 62:e02227–17.

43. Bouklas T, Alonso-Crisóstomo L, Székely T, Jr., Diago-Navarro E, Orner EP, Smith K, Munshi MA, Del Poeta M, Balázsi G, Fries BC. 2017. Generational distribution of a Candida glabrata population: Resilient old cells prevail, while younger cells dominate in the vulnerable host. PLOS Pathogens 13:e1006355.

44. Jain N, Cook E, Xess I, Hasan F, Fries D, Fries BC. 2009. Isolation and Characterization of Senescent *Cryptococcus neoformans* and Implications for Phenotypic Switching and Pathogenesis in Chronic Cryptococcosis. Eukaryotic Cell 8:858–866.

45. Walker LA, Munro CA. 2020. Caspofungin Induced Cell Wall Changes of Candida Species Influences Macrophage Interactions. Frontiers in Cellular and Infection Microbiology 10.

46. Kalem MC, Subbiah H, Leipheimer J, Glazier VE, Panepinto JC, Alspaugh JA. 2021. Puf4 Mediates Post-transcriptional Regulation of Cell Wall Biosynthesis and Caspofungin Resistance in Cryptococcus neoformans. mBio 12:e03225–20.

47. Garcia-Rubio R, Hernandez RY, Clear A, Healey KR, Shor E, Perlin DS. 2021. Critical Assessment of Cell Wall Integrity Factors Contributing to in vivo Echinocandin Tolerance and Resistance in Candida glabrata. Frontiers in microbiology 12:702779–702779.

48. Powell CD, Quain DE, Smart KA. 2003. Chitin scar breaks in aged Saccharomyces cerevisiae. Microbiology (Reading) 149:3129–3137.

49. Figueiredo ABC, Fonseca FL, Kuczera D, Conte FP, Arissawa M, Rodrigues ML. 2021. Monoclonal Antibodies against Cell Wall Chitooligomers as Accessory Tools for the Control of Cryptococcosis. Antimicrob Agents Chemother 65:e0118121.

50. 50. Lam WC, Upadhya R, Specht CA, Ragsdale AE, Hole CR, Levitz SM, Lodge JK. 2019. Chitosan Biosynthesis and Virulence in the Human Fungal Pathogen Cryptococcus gattii. mSphere 4.

51. Baker LG, Specht CA, Lodge JK. 2011. Cell wall chitosan is necessary for virulence in the opportunistic pathogen Cryptococcus neoformans. Eukaryot Cell 10:1264–8.

52. Upadhya R, Baker Lorina G, Lam Woei C, Specht Charles A, Donlin Maureen J, Lodge Jennifer K, Kronstad James W, Alspaugh JA, Lin X. Cryptococcus neoformans Cda1 and Its Chitin Deacetylase Activity Are Required for Fungal Pathogenesis. mBio 9:e02087–18.

53. Upadhya R, Lam WC, Hole CR, Parchment D, Lee CK, Specht CA, Levitz SM, Lodge JK. 2021. Cryptococcus neoformans Cda1 and Cda2 coordinate deacetylation of chitin during infection to control fungal virulence. The Cell Surface 7:100066.

54. Upadhya R, Lam WC, Maybruck B, Specht CA, Levitz SM, Lodge JK. 2016. Induction of Protective Immunity to Cryptococcal Infection in Mice by a Heat-Killed, Chitosan-Deficient Strain of Cryptococcus neoformans. mBio 7.

55. Nosanchuk JD, Stark RE, Casadevall A. 2015. Fungal Melanin: What do We Know About Structure? Frontiers in Microbiology 6.

56. van Duin D, Casadevall A, Nosanchuk JD. 2002. Melanization of Cryptococcus neoformans and Histoplasma capsulatum reduces their susceptibilities to amphotericin B and caspofungin. Antimicrob Agents Chemother 46:3394–400.

57. Chrissian C, Camacho E, Fu MS, Prados-Rosales R, Chatterjee S, Cordero RJB, Lodge JK, Casadevall A, Stark RE. 2020. Melanin deposition in two Cryptococcus species depends on cell-wall composition and flexibility. J Biol Chem 295:1815–1828.

58. Chrissian C, Lin CP, Camacho E, Casadevall A, Neiman AM, Stark RE. 2020. Unconventional Constituents and Shared Molecular Architecture of the Melanized Cell Wall of C. neoformans and Spore Wall of S. cerevisiae. J Fungi (Basel) 6.

59. Almeida-Paes R, Figueiredo-Carvalho MH, da Silva LB, Gerfen G, GR SA, Frases S, Zancopé-Oliveira RM, Nosanchuk JD. 2021. Candida glabrata produces a melanin-like pigment that protects against stress conditions encountered during parasitism. Future Microbiol 16:509–520.

60. Roy B, Jacobson A. 2013. The intimate relationships of mRNA decay and translation. Trends Genet 29:691–9.

61. Mukaremera L, Lee KK, Wagener J, Wiesner DL, Gow NAR, Nielsen K. 2018. Titan cell production in Cryptococcus neoformans reshapes the cell wall and capsule composition during infection. The Cell Surface 1:15–24.

62. Wiesner DL, Specht CA, Lee CK, Smith KD, Mukaremera L, Lee ST, Lee CG, Elias JA, Nielsen JN, Boulware DR, Bohjanen PR, Jenkins MK, Levitz SM, Nielsen K. 2015. Chitin Recognition via Chitotriosidase Promotes Pathologic Type-2 Helper T Cell Responses to Cryptococcal Infection. PLOS Pathogens 11:e1004701.

63. Lim J, Coates CJ, Seoane PI, Garelnabi M, Taylor-Smith LM, Monteith P, Macleod CL, Escaron CJ, Brown GD, Hall RA, May RC. 2018. Characterizing the Mechanisms of Nonopsonic Uptake of Cryptococci by Macrophages. J Immunol 200:3539–3546.

64. Gantner BN, Simmons RM, Underhill DM. 2005. Dectin-1 mediates macrophage recognition of Candida albicans yeast but not filaments. Embo j 24:1277–86.

65. Hasim S, Allison DP, Retterer ST, Hopke A, Wheeler RT, Doktycz MJ, Reynolds TB. 2016. β-(1,3)-Glucan Unmasking in Some Candida albicans Mutants Correlates with Increases in Cell Wall Surface Roughness and Decreases in Cell Wall Elasticity. Infection and immunity 85:e00601–16.

66. Wang ZA, Li LX, Doering TL. 2018. Unraveling synthesis of the cryptococcal cell wall and capsule. Glycobiology 28:719–730.

67. Jesus MD, Nicola AM, Chow SK, Lee IR, Nong S, Specht CA, Levitz SM, Casadevall A. 2010. Glucuronoxylomannan, galactoxylomannan, and mannoprotein occupy spatially separate and discrete regions in the capsule of Cryptococcus neoformans. Virulence 1:500–8.

68. Cordero RJB, Pontes B, Guimarães AJ, Martinez LR, Rivera J, Fries BC, Nimrichter L, Rodrigues ML, Viana NB, Casadevall A, Deepe GS. 2011. Chronological Aging Is Associated with Biophysical and Chemical Changes in the Capsule of Cryptococcus neoformans. Infection and Immunity 79:4990–5000.

69. 69. Reuwsaat JCV, Motta H, Garcia AWA, Vasconcelos CB, Marques BM, Oliveira NK, Rodrigues J, Ferrareze PAG, Frases S, Lopes W, Barcellos VA, Squizani ED, Horta JA, Schrank A, Rodrigues ML, Staats CC, Vainstein MH, Kmetzsch L. 2018. A Predicted Mannoprotein Participates in Cryptococcus gattii Capsular Structure. mSphere 3.

70. Bie X, Zhang S, Luo X, Qi RQ. 2019. Candida albicans cell wall mannoprotein synergizes with lipopolysaccharide to affect RAW264.7 proliferation, phagocytosis and apoptosis. Microb Pathog 131:98–105.

71. Ene IV, Walker LA, Schiavone M, Lee KK, Martin-Yken H, Dague E, Gow NAR, Munro CA, Brown AJP, Taylor JW. 2015. Cell Wall Remodeling Enzymes Modulate Fungal Cell Wall Elasticity and Osmotic Stress Resistance. mBio 6:e00986–15.

72. Bouklas T, Pechuan X, Goldman DL, Edelman B, Bergman A, Fries BC, Nielsen K, Berman J. 2013. Old Cryptococcus neoformans Cells Contribute to Virulence in Chronic Cryptococcosis. mBio 4:e00455–13.

73. Egilmez NK, Chen JB, Jazwinski SM. 1990. Preparation and partial characterization of old yeast cells. J Gerontol 45:B9–17.

74. Richards A, Gow NAR, Veses V. 2012. Identification of vacuole defects in fungi. Journal of Microbiological Methods 91:155–163.

75. Ding H, Caza M, Dong Y, Arif AA, Horianopoulos LC, Hu G, Johnson P, Kronstad JW, Deepe GS. 2018. *ATG* Genes Influence the Virulence of Cryptococcus neoformans through Contributions beyond Core Autophagy Functions. Infection and Immunity 86:e00069–18.

76. Fader CM, Colombo MI. 2009. Autophagy and multivesicular bodies: two closely related partners. Cell Death & Differentiation 16:70–78.

77. Park Y-D, Chen SH, Camacho E, Casadevall A, Williamson PR. 2020. Role of the ESCRT Pathway in Laccase Trafficking and Virulence of Cryptococcus neoformans. Infection and immunity 88:e00954–19.

78. Zhu X, Gibbons J, Garcia-Rivera J, Casadevall A, Williamson PR. 2001. Laccase of Cryptococcus neoformans is a cell wall-associated virulence factor. Infect Immun 69:5589–96.

79. Eisenman HC, Frases S, Nicola AM, Rodrigues ML, Casadevall A. 2009. Vesicle-associated melanization in Cryptococcus neoformans. Microbiology (Reading, England) 155:3860–3867.

80. Casadevall A, Nosanchuk JD, Williamson P, Rodrigues ML. 2009. Vesicular transport across the fungal cell wall. Trends Microbiol 17:158–62.

81. Rizzo J, Wong SSW, Gazi AD, Moyrand F, Chaze T, Commere P-H, Novault S, Matondo M, Péhau-Arnaudet G, Reis FCG, Vos M, Alves LR, May RC, Nimrichter L, Rodrigues ML, Aimanianda V, Janbon G. 2021. Cryptococcus extracellular vesicles properties and their use as vaccine platforms. Journal of extracellular vesicles 10:e12129–e12129.

82. Karkowska-Kuleta J, Kulig K, Karnas E, Zuba-Surma E, Woznicka O, Pyza E, Kuleta P, Osyczka A, Rapala-Kozik M, Kozik A. 2020. Characteristics of Extracellular Vesicles Released by the Pathogenic Yeast-Like Fungi Candida glabrata, Candida parapsilosis and Candida tropicalis. Cells 9:1722.

83. Zhao K, Bleackley M, Chisanga D, Gangoda L, Fonseka P, Liem M, Kalra H, Al Saffar H, Keerthikumar S, Ang C-S, Adda CG, Jiang L, Yap K, Poon IK, Lock P, Bulone V, Anderson M, Mathivanan S. 2019. Extracellular vesicles secreted by Saccharomyces cerevisiae are involved in cell wall remodelling. Communications Biology 2:305.

84. Albuquerque PC, Nakayasu ES, Rodrigues ML, Frases S, Casadevall A, Zancope-Oliveira RM, Almeida IC, Nosanchuk JD. 2008. Vesicular transport in Histoplasma capsulatum: an effective mechanism for trans-cell wall transfer of proteins and lipids in ascomycetes. Cell Microbiol 10:1695–710.

85. 85. Rizzo J, Chaze T, Miranda K, Roberson RW, Gorgette O, Nimrichter L, Matondo M, Latgé JP, Beauvais A, Rodrigues ML. 2020. Characterization of Extracellular Vesicles Produced by Aspergillus fumigatus Protoplasts. mSphere 5.

86. Lindstrom DL, Gottschling DE. 2009. The mother enrichment program: a genetic system for facile replicative life span analysis in Saccharomyces cerevisiae. Genetics 183:413–22, 1si-13si.

87. Wuttke D, de Magalhães JP. 2012. Osh6 links yeast vacuolar functions to lifespan extension and TOR. Cell Cycle 11:2419–2419.

88. Hughes AL, Gottschling DE. 2012. An early age increase in vacuolar pH limits mitochondrial function and lifespan in yeast. Nature 492:261–5.

89. Aufschnaiter A, Büttner S. 2019. The vacuolar shapes of ageing: From function to morphology. Biochimica et Biophysica Acta (BBA) - Molecular Cell Research 1866:957–970.

90. Rane HS, Hayek SR, Frye JE, Abeyta EL, Bernardo SM, Parra KJ, Lee SA. 2019. Candida albicans Pma1p Contributes to Growth, pH Homeostasis, and Hyphal Formation. Frontiers in Microbiology 10.

91. Martínez-Muñoz GA, Kane P. 2008. Vacuolar and plasma membrane proton pumps collaborate to achieve cytosolic pH homeostasis in yeast. The Journal of biological chemistry 283:20309–20319.

92. Bouklas T, Masone L, Fries BC. 2018. Differences in Sirtuin Regulation in Response to Calorie Restriction in Cryptococcus neoformans. J Fungi (Basel) 4.

93. Jain N, Bouklas T, Gupta A, Varshney AK, Orner EP, Fries BC. 2016. ALL2, a Homologue of ALL1, Has a Distinct Role in Regulating pH Homeostasis in the Pathogen Cryptococcus neoformans. Infect Immun 84:439–51.

94. Harrison TS, Chen J, Simons E, Levitz SM. 2002. Determination of the pH of the Cryptococcus neoformans vacuole. Med Mycol 40:329–32.

95. O’Meara TR, Holmer SM, Selvig K, Dietrich F, Alspaugh JA. 2013. Cryptococcus neoformans Rim101 Is Associated with Cell Wall Remodeling and Evasion of the Host Immune Responses. mBio 4:e00522–12.

96. Fuchs BB, Tegos GP, Hamblin MR, Mylonakis E. 2007. Susceptibility of *Cryptococcus neoformans* to Photodynamic Inactivation Is Associated with Cell Wall Integrity. Antimicrobial Agents and Chemotherapy 51:2929–2936.

97. García-Rivera J, Chang YC, Kwon-Chung KJ, Casadevall A. 2004. Cryptococcus neoformans CAP59 (or Cap59p) is involved in the extracellular trafficking of capsular glucuronoxylomannan. Eukaryot Cell 3:385–92.

98. Rizzo J, Colombo AC, Zamith-Miranda D, Silva VKA, Allegood JC, Casadevall A, Del Poeta M, Nosanchuk JD, Kronstad JW, Rodrigues ML. 2018. The putative flippase Apt1 is required for intracellular membrane architecture and biosynthesis of polysaccharide and lipids in Cryptococcus neoformans. Biochim Biophys Acta Mol Cell Res 1865:532–541.

